# An epigenetic basis of adaptive plasticity in *Drosophila melanogaster*

**DOI:** 10.1101/2022.10.11.511590

**Authors:** Abigail DiVito Evans, Regina A. Fairbanks, Paul Schmidt, Mia T. Levine

## Abstract

Fluctuating environments threaten fertility and viability. To better match the immediate, local environment, many organisms adopt alternative phenotypic states, a phenomenon called “phenotypic plasticity”. Local adaptation shapes phenotypic plasticity: natural populations that predictably encounter fluctuating environments tend to be more plastic than conspecific populations that encounter a constant environment. Despite pervasive evidence of such “adaptive phenotypic plasticity,” the evolution of the gene regulatory mechanisms underlying plasticity remains poorly understood. Here we test the hypothesis that environment-dependent phenotypic plasticity is mediated by epigenetic factors and that these epigenetic factors vary across naturally occurring genotypes. To test these hypotheses, we exploit the adaptive reproductive arrest of *Drosophila melanogaster* females, called diapause. Using an inbred line from a natural population with high diapause plasticity, we demonstrate that diapause is determined epigenetically: only a subset of genetically identical individuals enter diapause and this diapause plasticity is epigenetically transmitted for at least three generations. Upon screening a suite of epigenetic marks, we discovered that the active histone marks H3K4me3 and H3K36me1 are depleted in diapausing ovaries. Using ovary-specific knockdown of histone mark writers and erasers, we demonstrate that H3K4me3 and H3K36me1 depletion promotes diapause. Given that diapause is highly polygenic – distinct suites of alleles mediate diapause plasticity across distinct genotypes – we investigated the potential for genetic variation in diapause-determining epigenetic marks. Specifically, we asked if these histone marks were similarly depleted in diapause of a geographically distinct, comparatively less plastic genotype. We found evidence of genotypic divergence in both the gene expression program and histone mark abundance. This study reveals chromatin determinants of adaptive plasticity and suggests that these determinants are genotype-dependent, offering new insight into how organisms may exploit and evolve epigenetic mechanisms to persist in fluctuating environments.

## INTRODUCTION

Fluctuating environments threaten survival and reproduction in natural populations. The evolution of environment-dependent, phenotypic plasticity promotes the development of alternative phenotypes that better match the immediate, local environment. For example, seasonal snow cover triggers a coat color change from brown to white in the boreal snowshoe hare. Once the snow cover melts, the hare redevelops a brown coat (1). Similarly, limited resource availability triggers *Caenorhabditis elegans* juveniles to enter physiological arrest. Once resource availability improves, the juveniles resume development into adults (2, 3). Across a species’ range, the degree of environmental fluctuation may vary and selects for different degrees of phenotypic plasticity. Despite the clear relevance of such adaptive phenotypic plasticity in organismal and population responses to a changing climate, the molecular mechanisms that determine environment-induced plasticity are poorly understood (4–8).

Alternative plastic phenotypes are determined by coordinated up- and down-regulation of large swaths of the genome in response to changes in environmental conditions [reviewed in (5, 6)]. The molecular mechanisms that regulate alternative gene expression programs associated with phenotypic plasticity are largely unknown. In contrast, the gene regulatory mechanisms of cell fate plasticity are well-established. Epigenetic mechanisms such as DNA packaging into alternative “chromatin states” regulate cell fate plasticity by determining distinct gene expression programs and, ultimately, distinct cellular identities [(9–12) reviewed in (13–15)]. These alternative chromatin states include differential chemical modifications to either the DNA or the histone proteins that make up the nucleosome around which DNA wraps. The addition and removal of acetyl and methyl groups from histone tails can alter the transcriptional state of the underlying DNA and promote distinct cell fates in response to intrinsic developmental cues [(16, 17), reviewed in (18, 19)]. Intriguingly, extrinsic environmental cues can also alter DNA packaging into chromatin (20–26). Drought, temperature, salinity, and exposure to toxins alter the genome-wide distribution and abundance of acetyl and methyl groups on histone tails [reviewed in (27, 28)]. The observation that chromatin state is both environment-sensitive and a key determinant of cell fate during development raises the possibility that chromatin may mediate environment-sensitive phenotypic plasticity (29).

Consistent with this possibility, a handful of studies have established causal links between chromatin and phenotypic plasticity (29–34). Three of these studies probe chromatin-based regulation of reversible developmental arrest. To escape unfavorable environmental conditions, some organisms have evolved a state of dormancy in which development is suspended and senescence is slowed. Dormancy can occur at any developmental stage, from embryo to adult. In juvenile dormancy, a paused transition between developmental stages results in dramatic lifespan extensions. In adult dormancy, both somatic lifespan and reproductive lifespan are extended. While several groundbreaking studies have identified chromatin-based regulation of juvenile dormancy (32–34), the chromatin determinants of adult dormancy, and specifically the adaptive preservation of reproductive potential at this life stage, have not yet been explored. Moreover, we know virtually nothing about potentially adaptive genetic variation in the epigenetic mechanisms that promote and constrain phenotypic plasticity (35).

To investigate chromatin-based regulation of adaptive reproductive preservation in dormancy, we exploit the tractable model system, *Drosophila melanogaster. D. melanogaster* enters a form of adult dormancy called diapause in response to the cold temperatures and short days of oncoming winter [as defined in (36–39), but see (40)]. *Drosophila* diapause in females is characterized by extensive physiological changes that result in increased lipid storage, increased stress tolerance, increased lifespan extension, and suspended egg production. Suspended egg production is associated with global changes to the ovary transcriptome (41–46) and results in retention of nearly full reproductive potential following diapause (38, 43, 47, 48). Global changes to the ovary transcriptome under diapause implicates chromatin regulation, making diapause an ideal model to study the epigenetic determinants of reproductive dormancy.

*D. melanogaster* diapause is also an ideal model for investigating the evolution of gene regulatory mechanisms that mediate plasticity. Diapause plasticity is highly polygenic such that genetically distinct individuals have only partially overlapping suites of alleles that promote diapause (49). Diapause plasticity also varies adaptively (44, 50, 51). In populations from high latitudes with extreme winters, a higher proportion of females enter diapause under simulated winter conditions than females from low latitudes with mild winters (44, 50, 51). Similarly, a higher proportion of females enter diapause in populations collected immediately following winter than those collected in the late summer (51). This spatial- and temporal- variation in diapause plasticity, along with the observation that diapause plasticity is highly polygenic in *D. melanogaster*, makes this system ideal for probing how epigenetic determinants of plasticity vary across distinct genotypes.

Here we identify two epigenetic factors that regulate reproductive diapause through a mechanism distinct from those previously identified in juvenile diapause (32–34). We also show that these epigenetic determinants may vary across geographically distinct genotypes. These data provide new insight into how organisms exploit epigenetic mechanisms to persist in fluctuating environments, and how genetic variation may shape these epigenetic mechanisms.

## RESULTS

### Establishing a system to study epigenetic regulation of reproductive plasticity

To study the epigenetic determinants of reproductive lifespan extension under diapause, we established a system wherein epigenetic regulation, including chromatin-based gene regulation, could be isolated from the often-confounding effects of genotype, environment, and tissue heterogeneity. *D. melanogaster* diapause emerged as a compelling candidate system. This system allows us to control for genetic variation, to evaluate alternative developmental states in the same environment, and to ensure tissue homogeneity across alternative reproductive states.

To control for genotypic effects on epigenetics, we inbred an isofemale line from a temperate North American population (collected in Pennsylvania) by brother-sister mating for 10 generations. Under simulated winter conditions (Fig. 1A), 87.9% of inbred females enter diapause (Fig. 1B). Importantly, the incidence of diapause in this inbred line does not differ from the isofemale line from which it was derived (χ^2^ test, p = 0.49), suggesting that residual, segregating genetic variation alone does not account for the observed degree of plasticity. Incomplete diapause penetrance, where most females arrest but some (12.1%) remain persistently reproductive upon exposure to simulated winter conditions, allows us to control for environment in addition to genotype: we can compare the chromatin state of inbred diapausing and persistently reproductive individuals in the same environment. Finally, to control for tissue heterogeneity in all experiments that compared arrested and persistently reproductive ovaries, we isolated ovary stages 1-7 (see Methods). Stages 1-7 are those represented in diapause (Fig. 1B). This careful exclusion of development beyond stage 7 allowed us to control for cell type composition between arrested and persistently reproductive ovaries.

**Figure 1.**
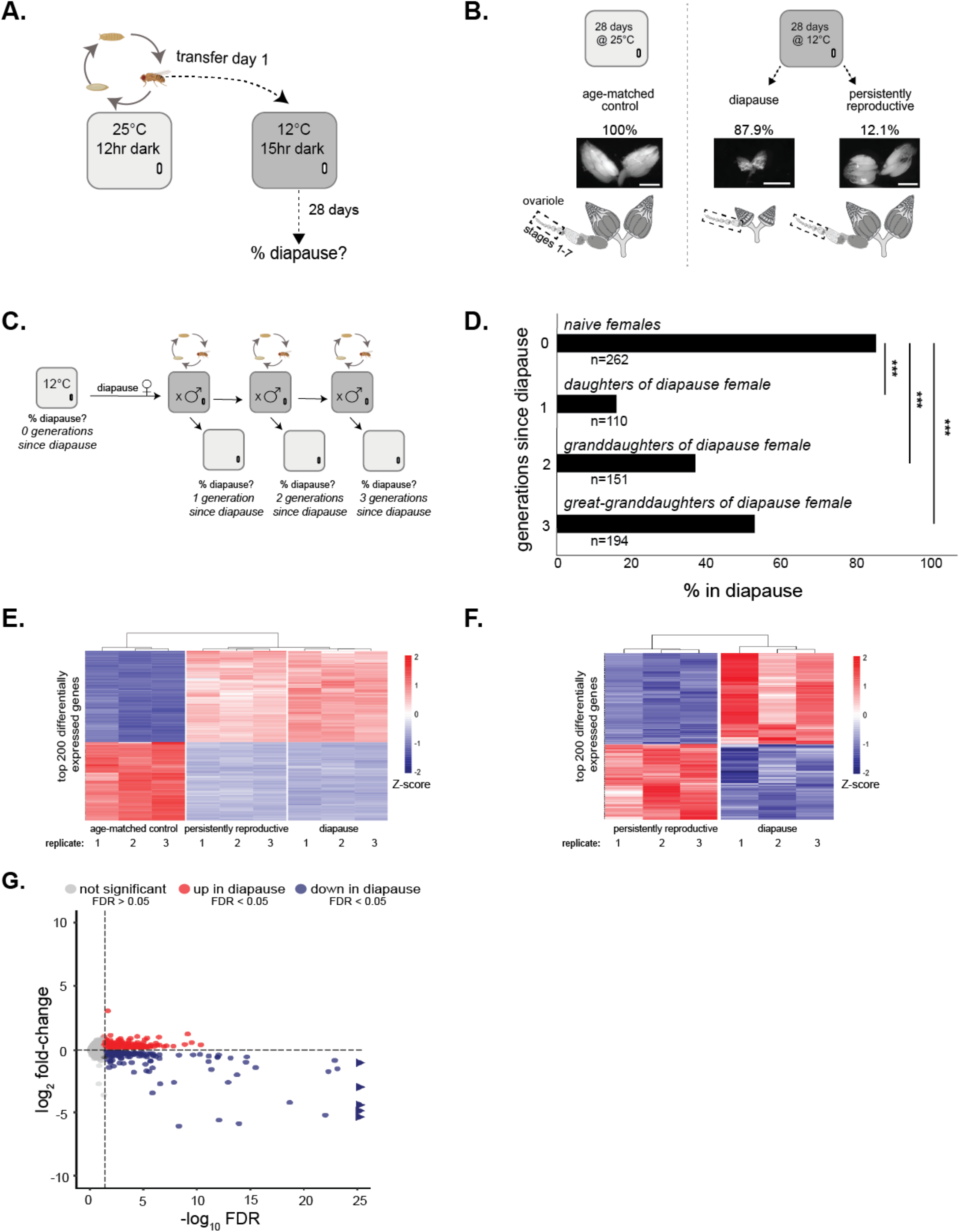
Establishing a system to study epigenetic regulation of adaptive phenotypic plasticity. (A) Diagram of diapause assay design. Flies are reared at 25°C under a 12-hour dark/light cycle in an incubator (light gray). Females are then transferred to an incubator set to 12°C under a 15-hour dark, 9-hour light cycle (dark gray, simulated winter conditions). Females are maintained in simulated winter conditions for 28 days before ovaries are assessed for diapause (% of females with arrested ovaries, i.e., “diapause plasticity”). (B) Degree of plasticity in age-matched control at 25°C (left) and diapause and persistently reproductive at 12°C (right). Ovaries from age-matched control females, diapause females, and persistently reproductive females and cartoons (below) representing the ovaries with a separated single ovariole, the basic unit of egg production in the *Drosophila* ovary. Dotted box indicates stages 1-7 used for all RNA and protein assays (note that the arrested ovary has only stages 1-7, created using Biorender.com). Scale bars = 0.5 mm (C) Diagram of transgenerational assay design. After maintenance under simulated winter conditions for 28 days (dark gray incubator, see above), females are assayed for diapause using a non-destructive method (see Methods). The females previously in diapause (“naïve females”) are then crossed to males at 25°C. Virgin females from this cross (“daughters of diapause female”) are placed either into a 12°C incubator to assess diapause plasticity or into a vial with males at 25°C to generate the granddaughters of diapause females. This process is repeated with these granddaughters and the great-granddaughters of diapause females. (D) Diapause plasticity of daughters, granddaughters, and great-granddaughters of females who underwent diapause. χ^2^, *** p<0.001. (E) Heatmap of the top 200 differentially expressed genes (by FDR) between age-matched control, diapausing, and persistently reproductive ovaries. Blue-red gradient depicts the Z-score of each gene. Red corresponds to upregulated genes and blue corresponds to downregulated genes. (F) Heatmap of top 200 differentially expressed genes (by FDR) between diapause and persistently reproductive ovaries. Blue-red gradient depicts the Z-score of each gene. Red corresponds to upregulated genes and blue corresponds to downregulated genes. (G) Volcano plot showing differential gene expression across diapausing and persistently reproductive ovaries. Triangles represent genes with −log_10_ FDR > 25.

The observation of environment-induced alternative phenotypic states across individuals of a single genotype implicates epigenetic regulation. Another hallmark of epigenetic regulation is transgenerational transmission of parental environmental conditions to offspring (52–55). To probe the possibility that parental diapause is transmitted to offspring, we assayed diapause plasticity (the proportion of females that enter diapause) of daughters, granddaughters, and great-granddaughters of inbred females that had undergone diapause (Fig. 1C). We first subjected a cohort of females to winter conditions (12°C, Fig. 1C “naïve females”). We allowed the subset of females that had entered diapause to mate and reproduce at 25°C. We assayed one cohort of their daughters for diapause plasticity (Fig. 1C “daughters”, one generation since diapause) and allowed a second cohort to mate and reproduce at 25°C. We repeated this process until the great-granddaughter generation (Fig. 1C “great granddaughters”, three generations since diapause). We note that, under this protocol, the *propagated* daughters, granddaughters, and great-granddaughters are never exposed to winter conditions. A subset of these progeny was sampled to determine diapause propensity and then discarded. These experiments revealed that diapause entry in mothers reduces the proportion of daughters in diapause (p<2.2×10^-16^), granddaughters (p<2.2×10^-16^), and great-granddaughters (p=4.6×10^-8^), Fig. 1D). Moreover, this transgenerational effect decreases with each generation removed from the initial incidence of diapause. The dilution of the transgenerational effect suggests dilution of an epigenetic signal through generations [(56), reviewed in (57)]. Such transgenerational effects in an inbred line further implicate a role for epigenetic regulation of diapause plasticity.

A classic readout of epigenetically regulated, alternative phenotypic fates is alternative gene expression programs across genetically identical individuals (58). To profile gene expression across the two reproductive fates, we performed RNA-seq on both arrested and persistently reproductive ovaries from females maintained under simulated winter conditions for 28 days (see Methods). We also performed RNA-seq on ovaries from age-matched control females maintained at 25°C for 28 days. We prepared RNA from exclusively stages 1-7 in both arrested and reproductive ovaries to ensure tissue homogeneity between samples (Fig. 1B).

RNA-seq revealed distinct gene expression profiles of age-matched control ovaries (25°C), arrested ovaries (12°C), and persistently reproductive ovaries (12°C). Consistent with the well-documented, pervasive effects of temperature alone on gene expression (59, 60), the 200 most differentially expressed genes (by false discovery rate, “FDR”) are differentially expressed between age-matched control ovaries at 25°C and ovaries at 12°C (Fig. 1E, Fig. S1A); however, within the 12°C treatment, diapausing and persistently reproductive ovaries have distinct gene expression programs (Fig. 1F, Fig. S1B). More genes are down-regulated than up-regulated in diapausing compared to persistently reproductive ovaries, and more down-regulated genes have log2-fold change greater than two (Fig. 1G). Nevertheless, the significant upregulation of hundreds of genes in diapause suggests that *D. melanogaster* diapause is not simply a generalized shut-down of gene expression but instead an actively regulated state [see also (46)]. This differential gene expression between diapause and persistently reproductive ovaries at 12°C, combined with the transgenerational effect of diapause, suggests that epigenetic factors mediate reproductive arrest in the ovary.

### Epigenetic marks H3K4me3 and H3K36me1 regulate diapause plasticity

Epigenetic regulation depends in part on chromatin modifications [reviewed in (61)]. The basic unit of chromatin is the nucleosome, the octameric complex of histone proteins around which DNA wraps (62). Residues on the tails of histones can be post-translationally modified, primarily by the addition or removal of acetyl and methyl groups (63). These histone marks can alter the transcriptional activity of the underlying DNA [reviewed in (64)]. To identify histone marks associated with diapause plasticity, we prepared lysate from arrested ovaries and persistently reproductive ovaries (stages 1-7 only) and screened six, highly abundant histone H3 modifications (65). Given the downregulation of most genes in diapausing ovaries (Fig. 1G), we predicted either an excess of repressive marks or the depletion of active marks. The screen revealed that repressive marks H3K27me3 and H3K9me3, as well as active marks H3K27ac and H3K9ac, did not differ in abundance across diapausing and persistently reproductive ovaries (Fig. 2A, Fig. S2). In contrast, active marks H3K4me3 and H3K36me1 were depleted in diapause (Fig. 2A, Fig. S2).

**Figure 2.**
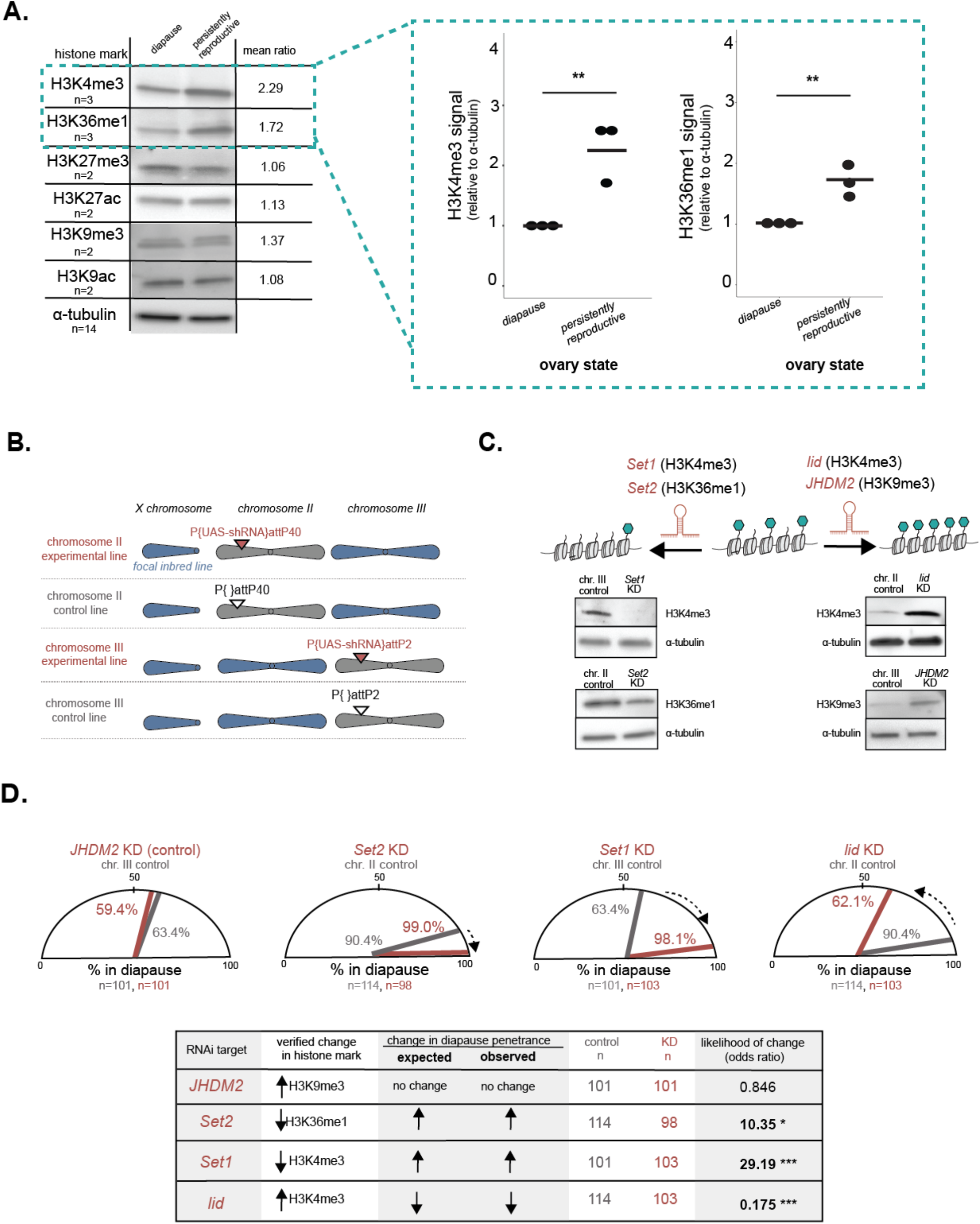
H3K4me3 and H3K36me1 regulate diapause plasticity. (A) Representative western blots showing histone mark abundance in diapausing and persistently reproductive ovaries. Quantification of H3K4me3 and H3K36me1 signal relative to a-tubulin loading control for three biological replicates (right). t-test, **p < 0.01. (B) Genotypes of experimental and control lines in histone mark manipulation experiment. Blue chromosomes represent chromosomes from the focal inbred line. Experimental genotypes encode a construct with a UAS promoter that drives a small hairpin RNA (“P{UAS-shRNA}”). These constructs are inserted into a chromosome-specific attP site (pink triangle). Control line attP sites lack the inserted construct (white triangle). All lines are crossed to the same ovary-specific driver. (C) Expected change of histone mark abundance after RNAi against histone mark writers and erasers (above) and western blots validating histone mark depletion or enrichment (below). “chr.” = chromosome. (D) Diapause plasticity of experimental (pink) and control (gray) genotypes. Expected direction of change shown with a black dotted arrow. FET, * p < 0.05, *** p < 0.001. “chr.” = chromosome. Note that we ruled out the possibility that diapause-independent effects of knockdown of *Set1* and *Set2* on ovary development confounded these results (see Methods and Fig. S3B)

The depletion of active marks in diapausing ovaries could be a byproduct of the overall downregulation of genes in diapause or could reflect a causal role of histone marks in determining diapause plasticity. To test the prediction that the abundance of these marks in the ovary affect reproductive plasticity, we manipulated histone mark abundance. Using the Gal4/UAS system (see Methods), we expressed in the ovary short hairpin RNAs (shRNAs) that knock down transcripts of enzymes that deposit or remove these histone marks (Fig. 2B,C). Given that diapause plasticity is well-known to vary across genotypes (42, 45, 66, 67), we strictly controlled the genetic background of the experimental and control flies. To generate the experimental lines, we introduced chromosomes carrying the focal shRNA construct, integrated into an attP landing site, into the inbred line from Pennsylvania described above (Fig. 2B). To generate the control lines, we introduced chromosomes that have the same attP landing site, but *lack* the shRNA construct, into the same inbred line (Fig. 2B). We crossed these experimental and control lines to a driver line that directs expression of the shRNA in the ovary.

We manipulated the abundance of three histone marks in the ovary and assayed diapause plasticity (the proportion of females with arrested ovaries) using these rigorously controlled genotypes. First, we manipulated a “control” histone mark, H3K9me3, which did not vary between diapausing and reproductive ovaries (Fig. 2A, Fig. S2). Specifically, we knocked down *JHDM2* (Fig. S3A), an enzyme that demethylates H3K9. As expected, *JHDM2* knockdown elevated H3K9me3 (Fig. 2C) but had no effect on diapause plasticity (Fig. 2D, odds ratio=0.846, p>0.05). Next, we manipulated H3K36me1 and H3K4me3, two histone marks depleted in arrested ovaries (Fig. 2A). We predicted that experimental depletion of these marks would increase diapause plasticity, while experimental enrichment would decrease diapause plasticity. This is exactly what we observed. To deplete H3K36me1, we knocked down *Set2*, which encodes an enzyme that methylates H3K36 (Fig. 2C, Fig. S3A). Indeed, H3K36me1 depletion increased diapause plasticity (Fig. 2D, odds ratio=10.35, p<0.05). Similarly, we depleted H3K4me3 by knocking down *Set1*, which encodes an enzyme that methylates H3K4 (Fig. 2C, Fig. S3A) and again observed increased diapause plasticity (Fig. 2D, odds ratio=29.19, p<0.0001). We then experimentally *enriched* H3K4me3 by knocking down *lid*, which encodes an enzyme that removes H3K4 methylation (Fig. 2C, Fig. S3A). As predicted, this opposing manipulation decreased diapause plasticity (Fig. 2D, odds ratio=0.175, p<0.0001). Observing this opposing effect of decreased diapause blunted our concern that active mark depletion simply blocks ovary development beyond stage 7. Furthermore, our observation of many persistently reproductive ovaries upon depletion of both H3K36me1 and H3K4me3 in the context of the diapause plasticity-increasing transgenerational effect also rejects the possibility that compromised ovary development confounds our results (Figure S3B, see Methods). Together, these data suggest that H3K36me1 and H3K4me3 depletion, but not H3K9me3 elevation, promotes diapause plasticity.

### Diapause-associated chromatin state and gene expression are genotype-specific

Diapause plasticity in *D. melanogaster* is a highly polygenic trait that varies both geographically and seasonally, as described above (44, 49–51). Because diapause is determined by variation at hundreds of genes, geographically distinct populations share only partially overlapping alleles that promote (or constrain) diapause plasticity. This distinct genetic architecture predicts distinct transcriptional programs across natural populations and raises the possibility that distinct epigenetic mechanisms contribute to diapause plasticity across distinct genotypes. To explore this possibility, we inbred an additional line collected from subtropical Florida, a region with mild winters. As expected, this inbred line has low diapause plasticity (14.9% diapause, Fig. 3A). Henceforth, we refer to the temperate inbred line described above as “High Plasticity” or “HP” (87.9% diapause), and the subtropical inbred line as “Low Plasticity” or “LP”.

**Figure 3.**
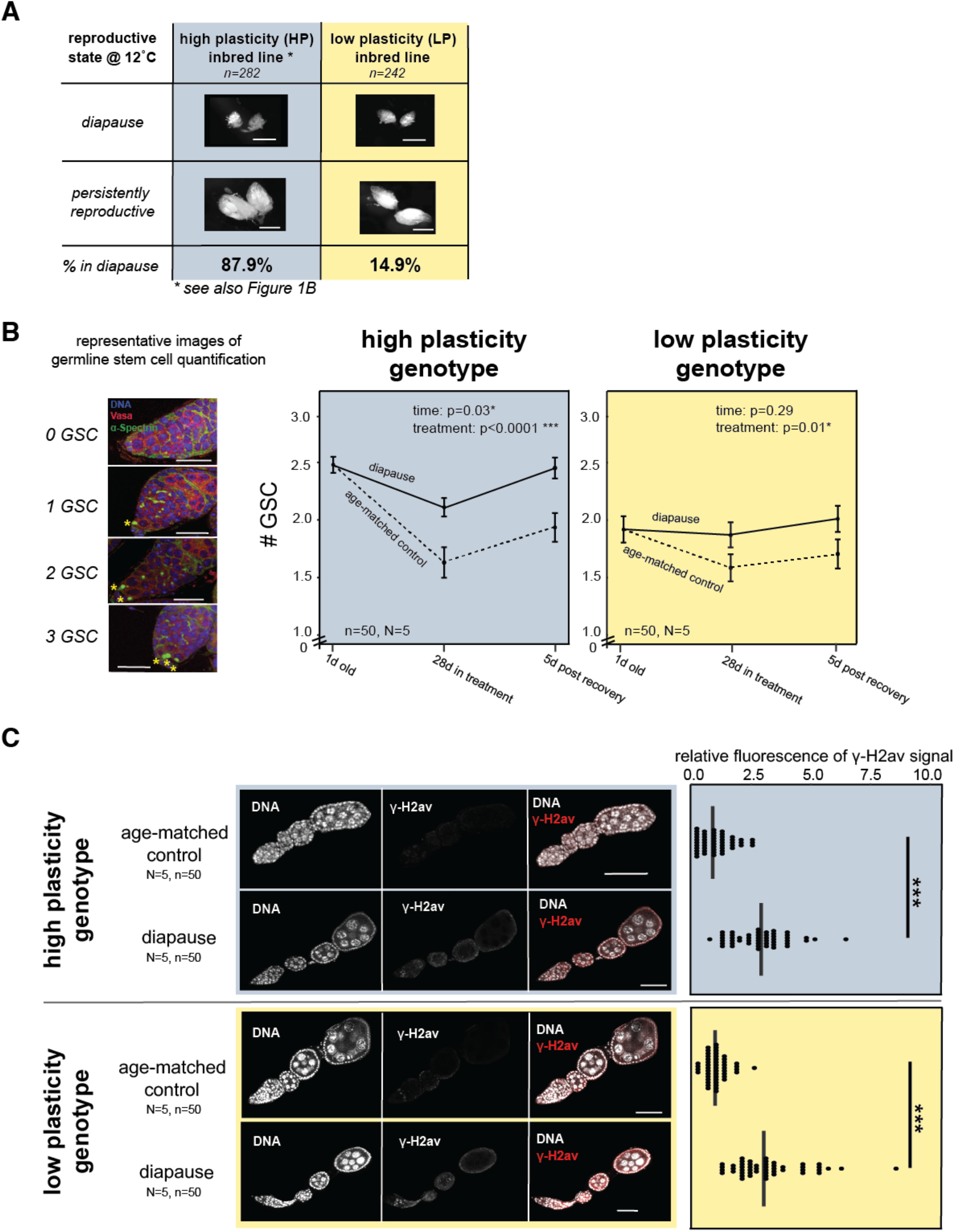
Assessing the diagnostic features of diapause in a low plasticity line. (A) Representative images of diapausing and persistently reproductive ovaries of high plasticity (HP, blue) and low plasticity (LP, yellow) inbred lines as well as the degree of plasticity (note: HP % diapause reported in Figure 1B). Scale bar = 0.5 mm. (B) Representative images of germaria with 0, 1, 2, and 3 germline stem cells (“GSC,” designated by *, scale bar = 10μm) stained with DAPI (blue), anti-Vasa (red) and anti-a-Spectrin (green). Average number of germline stem cells at one day old, after 28 days of treatment (25°C or diapause), and after five days post-treatment at 25°C for diapausing and age-matched control ovaries of HP and LP females (scale bar = 25μm, d=”day”, 2-way ANOVA with fixed effects = timepoint, treatment, error bars = SEM). (C) γ-H2av signal in age-matched control and diapausing ovaries of HP and LP females and quantification. Mann-Whitney U test, *** p<0.001, scale bar = 50μm.

Given that the LP line is largely insensitive to the simulated winter conditions, we first determined whether diapause in the LP line is a *bona fide* alternative developmental state or instead a generalized stress response to simulated winter conditions [see (68)]. The *D. melanogaster* female’s generalized stress response to unfavorable environmental conditions, like starvation or predator exposure, manifests superficially as arrested ovary development (69–72). However, generalized stress response in the ovaries is both cell biologically and functionally distinct from diapause (48). Previous studies have demonstrated that diapause preserves fertility and germline stem cell number compared to age-matched controls, while stress does not (48). Diapausing ovaries also accumulate the double-strand break marker, γ-H2av, due to the persistence of egg chambers in extended arrest (48). We found that despite low responsiveness to simulated winter conditions, the LP line, like the HP line, preserves germline stem cell number in diapause compared to age-matched controls (Fig. 3B). Furthermore, LP and HP diapausing ovaries are similarly enriched for γ-H2av compared to age-matched controls (Fig. 3C). Diapause also preserves fertility in the LP line compared to age-matched controls (Fig. S4). These data suggest that the LP line enters a true diapause state in response to simulated winter conditions.

To investigate whether the epigenetic mechanisms mediating plasticity are distinct across diverged genotypes, we first asked if genetic variation in diapause plasticity manifests as transcriptional variation. We conducted RNA-seq on diapausing and persistently reproductive ovaries from the LP line and compared the statistically significant differential gene expression to that of the HP line (Fig. S5, Fig. 1E). To isolate those genes that are up- and down-regulated specifically in diapause, we normalized the list of genes that were differentially expressed between diapausing and persistently reproductive ovaries (both at 12°C) to gene expression of age-matched control ovaries (25°C). Specifically, we included in downstream analyses only those genes that were differentially expressed between diapausing and persistently reproductive ovaries *and* between diapausing and age-matched control ovaries in a given genotype, removing genotype-specific expression that was independent of diapause. We compared this reduced list of diapause-specific genes across the two genotypes (433 in HP, 606 in LP) and determined which genes were differentially expressed in both HP and LP (“genotype-independent”), differentially expressed only in the HP line (“HP-specific”), or differentially expressed only in the LP line (“LP-specific”). While many genes that are up- or down-regulated in diapause are shared across the HP and LP lines (201 genes, Fig. 4A), most differentially expressed genes are genotype-dependent (611 genes, Fig. 4A). Consistent with the known polygenic basis of diapause plasticity, these results suggest that HP and LP diapause plasticity are associated with only partially overlapping transcriptional programs.

**Figure 4.**
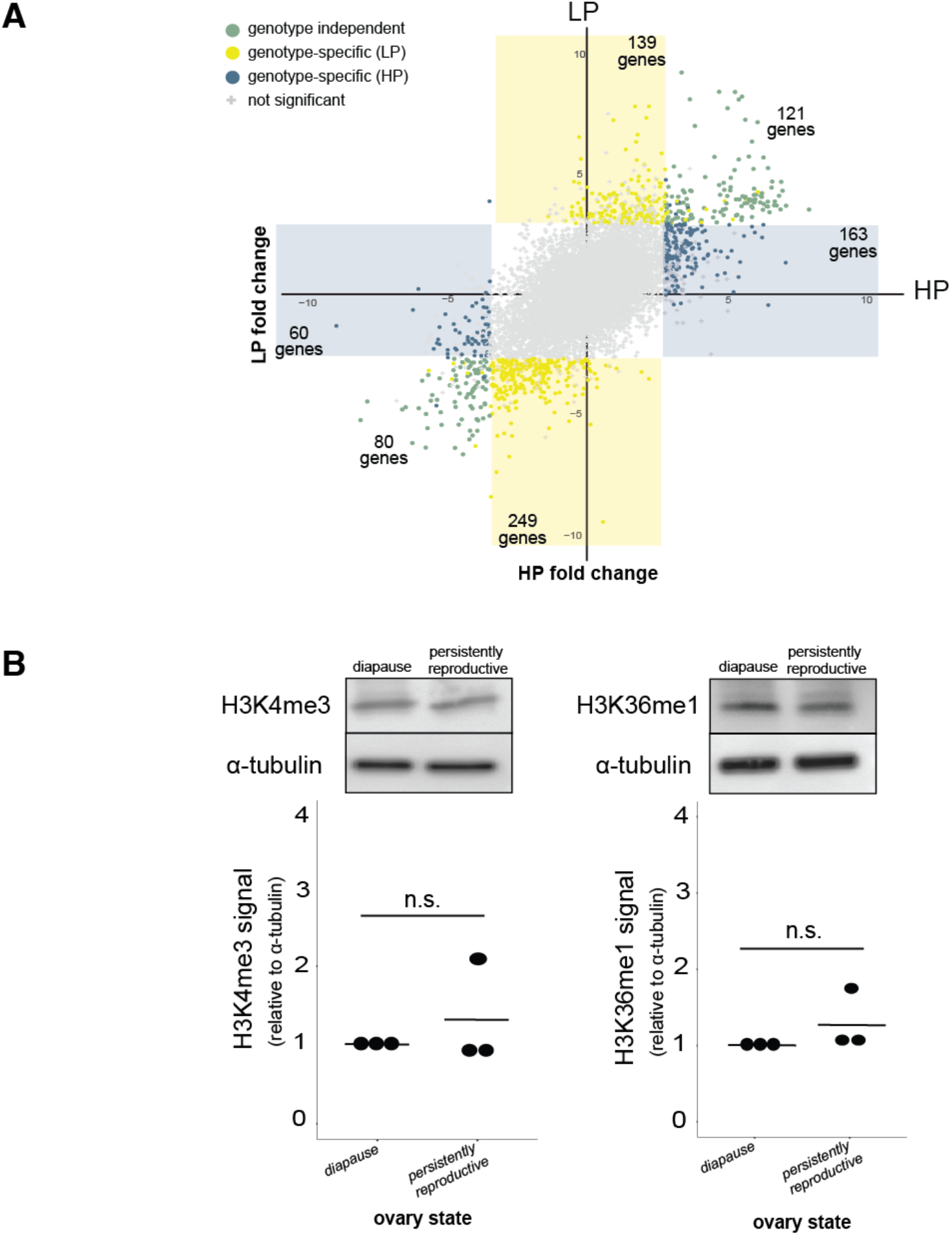
Diapause-associated gene expression and diapause-associated chromatin state are genotype-specific. (A) Differential gene expression across diapausing and persistently reproductive ovaries in HP and LP genotypes. Gray points represent genes that are not significantly differentially expressed in either genotype. Green points represent genes that are differentially expressed in both genotypes (FDR < 0.05). Blue points represent genes that are differentially expressed in the HP genotype only (FDR < 0.05). Yellow points represent genes that are differentially expressed in the LP genotype only (FDR < 0.05). (B) Representative western blots showing the abundance of H3K4me3 and H3K36me1 in diapausing and persistently reproductive ovaries from the LP inbred genotype females (above). Quantifications of H3K4me3 and H3K36me1 signal relative to a-tubulin loading control for three replicates (below). t-test, n.s. = p>0.05, (compare to Figure 2A).

We predicted that the genes up- or down-regulated in diapausing ovaries in only one genotype (“genotype-dependent”) may regulate pathways that promote reproductive arrest common to both genotypes. Consistent with this prediction, we found evidence that genotype-dependent gene expression in HP and LP diapause converges on common biological processes. For example, two metabolic pathways involved in ATP synthesis (“The citric acid (TCA) cycle and respiratory electron transport” and “Respiratory electron transport, ATP synthesis by chemiosmotic coupling, and heat production by uncoupling proteins”) are enriched in both HP-specific and LP-specific genes up-regulated in diapause (Table 1, highlighted in green). Moreover, twelve pathways are enriched for *both* genotype-independent genes and genotype-*dependent* genes upregulated in diapause (Table 1, highlighted in gray). This finding suggests that some pathways are utilized by both genotypes via the expression of overlapping genes *and* non-overlapping genes. There were no significant pathways overrepresented for genes downregulated in HP-dependent genes and only a single pathway overrepresented for genotype-independent downregulated genes; consequently, common pathways could not be detected for down-regulated genes. These results suggest that diapause in both genotypes depends on the activation of common pathways despite pervasive genotype-dependent gene expression.

**Table 1.**
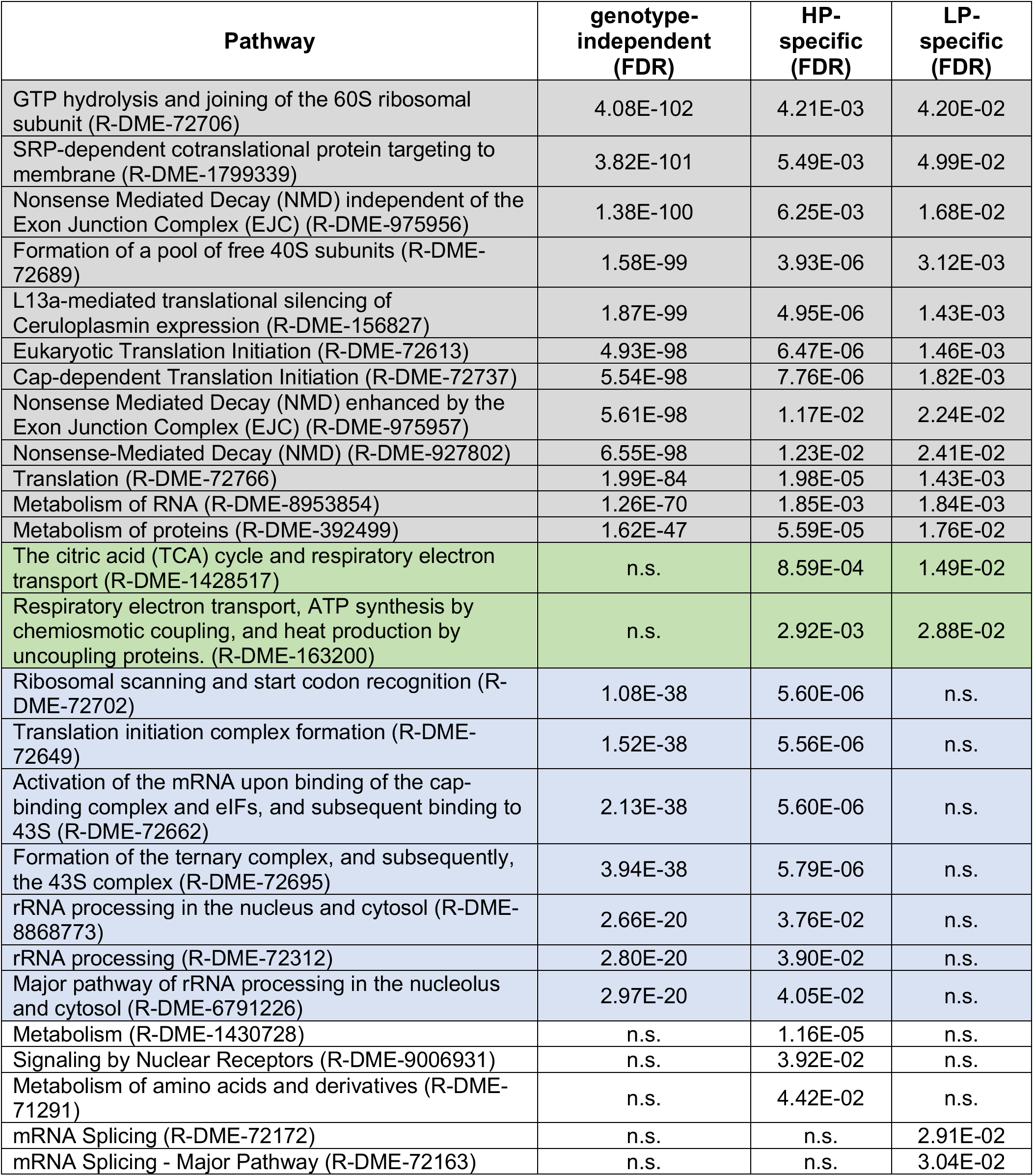
Pathway enrichment for genes up-regulated in diapause. Gray pathways are significant in all categories (genotype-independent, HP-specific and LP-specific genes). Green pathways are significant in both HP-specific and LP-specific genes. Blue pathways are significant in both genotype-independent and HP-specific genes. No pathways are significant in both genotype-independent and LP-specific genes.

Genotype-dependent gene expression raises two alternative hypotheses that could explain distinct epigenetic regulation of diapause across distinct genotypes. Diapause plasticity in the HP and LP lines could be regulated by distinct epigenetic marks at the distinct sets of genes associated with diapause. Alternatively, the same epigenetic mark, like H3K4me3, could be depleted at distinct sets of genes across the two genotypes. To evaluate these alternative hypotheses, we tried a genome-wide chromatin profiling approach that requires minimal sample material [CUT&RUN, (73)]. Unfortunately, attempts to profile the minimal tissue from stages 1-7 of the diapausing and persistently reproductive ovaries were unsuccessful. We therefore turned to histone mark abundance to determine whether distinct chromatin states underlie diapause in distinct genotypes. Similar to the HP line, H3K9ac, H3K9me3, H3K27ac and H3K27me3 abundance did not differ between diapausing and persistently reproductive ovaries of the LP line (Fig. S2). However, unlike the H3K4me3 and H3K36me1 depletion that we observed in diapausing ovaries of the HP line, these two marks were invariant across the two reproductive states in the LP line (Fig. 4B). These data are consistent with distinct chromatin states of diapausing ovaries across the HP and LP lines. These data also highlight the idea that global downregulation of the genome during diapause is not inevitably associated with loss of a pervasive, active histone mark. Future work will identify the epigenetic factors that determine gene expression and diapause plasticity in the LP line.

## DISCUSSION

Here we describe a new, tractable system for studying genetic variation in epigenetic regulation of adaptive phenotypic plasticity. Importantly, this model system controls for the confounding effects of genetic variation, environment, and tissue heterogeneity on chromatin packaging, allowing us to isolate functional links between chromatin modifications and phenotypic state. We discovered that environment-dependent reproductive arrest in *D. melanogaster* is mediated by at least two histone marks, H3K4me3 and H3K36me1. We also found that this epigenetic mechanism may be shaped by genetic variation – these two marks were depleted in diapausing ovaries of a temperate, high plasticity genotype but were similarly abundant across diapausing and reproductive ovaries in a subtropical, low plasticity genotype. These data raise the possibility of a distinct epigenetic basis of diapause plasticity across distinct genotypes.

Previous studies in non-model or emerging model systems have demonstrated compelling causal links between chromatin and diapause. Two major chromatin silencing pathways emerged from these studies: DNA methylation in *Nasonia* wasps (34) and H3K27me3 in the Cotton bollworm moth (33) and the Turquoise killifish (32). *D. melanogaster* has minimal DNA methylation (74); consequently, we focused on histone marks like H3K27me3. Surprisingly, the screen of histone marks revealed no difference in abundance of H3K27me3 between arrested and persistently reproductive ovaries [as was found in bollworm pupa (33)], and we did not detect differential expression of enzymes that write, read, or erase H3K27me3 [as was found in killifish embryos (32)]. These observations implicated a distinct chromatin mechanism regulating *D. melanogaster* reproductive diapause. Our focus on the ovary may account for this difference. Previous studies probed paused developmental transitions during juvenile phases, either embryonic or larval, rather than the adult reproductive tissues. Consistently, H3K27me3 is a classic regulator of developmental fate (75). It is also possible that H3K27me3, along with H3K4me3 and H3K36me1, regulates diapause plasticity but gross H3K27me3 abundance does not vary between reproductive states.

The discovery that H3K4me3 depletion promotes reproductive diapause is reminiscent of earlier studies of somatic aging. From yeast to mammals, aging is associated with a general increase in active chromatin marks like H3K4me3 and an overall increase in transcription [reviewed in (76)]. In *Drosophila*, H3K4me3 depletion extends lifespan (77). Like these studies of somatic aging, we found not only that H3K4me3 depletion promotes reproductive diapause but also that diapause is associated with a genome-wide decrease in gene expression in the high plastic (HP) genotype. These results suggest that the chromatin state of the diapausing HP ovary mirrors that of young somatic tissues. Given that the chromatin determinants of age-dependent reproductive decline (78) are not nearly as well explored as the determinants of somatic aging [reviewed in (76, 79)], this system offers a new foothold for understanding how a youthful chromatin state contributes to the preservation of reproductive potential.

Chromatin mediates not only development and aging within a generation but also mediates epigenetic information transfer between generations [reviewed in (80, 81)]. Our discovery of distinct epigenetic states between arrested and persistently reproductive ovaries raised the possibility that information about the maternal environment is transferred to the next generation. Consistent with diapause-induced transmission of epigenetic information, we found that diapause entry reduces subsequent diapause plasticity of genetically identical daughters, granddaughters, and great-granddaughters. Such transgenerational memory requires a DNA sequence-independent “message” to be passed from parent to offspring through the germline. The identity of this message remains mysterious; however, we speculate that either alternative chromatin packaging or specific transcripts, possibly small RNA(s) (82, 83), of diapausing ovaries transmits this heritable epigenetic information. The discovery of transgenerational information transfer from mother to daughter – in a model system – puts forward *D. melanogaster* diapause as a powerful new resource for studying epigenetic inheritance.

Our discovery of epigenetic regulation of a highly polygenic and adaptively varying trait offered us the unique opportunity to explore the effect of genetic variation on epigenetic regulation. This study brings together the historically distinct areas of epigenetic regulation and genetic variation in plasticity. We uncovered evidence of genetic variation of epigenetic marks causally linked to diapause plasticity. Unfortunately, the sensitivity of diapause plasticity to genetic background precluded us from using RNAi to test directly the hypothesis that the low plasticity, subtropical derived line is insensitive to H3K4me3 and H3K36me1 manipulation. Furthermore, our attempts to conduct genome-wide chromatin profiling across our two genotypes were unsuccessful. Innovation in genome-wide histone mark profiling on minimal tissue will allow us to further probe the possibility that diapause in the ovary is regulated by distinct epigenetic mechanisms in distinct genotypes.

Genotypic variation in the epigenetic regulation of diapause raises the possibility that distinct epigenetic factors are positively selected to confer distinct degrees of reproductive arrest. Intriguingly, genes encoding many epigenetic factors exhibit latitudinal clines, including enzymes that methylate and demethylate H3K4 and H3K36 (84–86). Future research comparing the chromatin regulation of diapause in many distinct genotypes from various geographic regions and seasonal timepoints may uncover spatial and temporal variation in epigenetic regulation of diapause. This system is now well-positioned to offer the first glimpses of how adaptive evolution shapes epigenetic mechanisms underlying adaptive phenotypic plasticity. Understanding these evolutionary processes is vital: genetic variation of epigenetic regulation likely shapes how natural populations respond to the extreme seasonal environments arising from ongoing climate change (87).

## MATERIALS AND METHODS

### *Drosophila* stocks and culturing

We constructed the High Plasticity (“HP”) inbred line from an isofemale line collected from Media, Pennsylvania in July, 2018. To inbreed the line, we mated one brother and one sister each generation for 10 generations. We similarly constructed the Low Plasticity (“LP”) line from an isofemale line collected in Miami, Florida in July 2018. We maintained stocks at 25°C in 12-hour light/dark cycles on standard molasses food.

### Diapause assay

To assay diapause, we subjected 0-6 hour-old virgin females to 12°C and a short-day light cycle (9 hours of light, 15 hours of darkness) for 28 days. To assay whether a female was in diapause, we dissected out the ovaries and determined the latest developmental stage of the ovary as defined in Saunders *et al*. [1989, (41)]. Specifically, we designated females as undergoing diapause if they lacked vitellogenic egg chambers (stage 8 or later, Fig. 1B). We designated females as “persistently reproductive” if both ovaries had one or more egg chambers at stage 8 or later. We excluded from experiments those rare females (<1%) whose ovaries fell between these categories.

### Transgenerational assay

To assess transgenerational effects of diapause, we induced diapause as described above but determined the reproductive state of females without dissection using a modification of the “Bellymount” protocol described in (88). We positioned females between two cover slips with a drop of 50% glycerol and determined the reproductive state by visualizing through the abdomen the presence or absence of egg chambers at stage 8 or later. We then crossed females in diapause (“naïve females”, Fig. 1C,D) to males from the same inbred line at 25°C. We placed virgin female offspring from this cross either into a 12°C incubator (“daughters of diapause females,” Fig. 1C,D) to assess diapause rate or into a vial with males at 25°C to generate granddaughters of the generation 0 diapause females. We repeated this process with these granddaughters as well as the great-granddaughters of the generation 0 diapause (“naïve”) females. We compared diapause plasticity of generation 0 to that of daughters, granddaughters, and great-granddaughters using χ^2^ test.

### RNA-sequencing and analysis

To define the gene expression associated with diapause while controlling for genotype, temperature, and age, we induced diapause as described above. We kept age-matched control flies as virgins in an incubator set to 25°C and a long day light cycle (12 hours of light, 12 hours of dark) for 28 days. We flipped these control females onto fresh food every 7 days due to accelerated mold growth on the food at 25°C. We isolated equivalent egg chamber stages from ovaries across reproductive states. For persistently reproductive ovaries and age-matched control ovaries, we dissected off accessory structures and then isolated by microdissection ovary stages 1-7 only (Fig. 1B). For arrested ovaries, we removed the accessory structures only. We prepared total RNA (Mirvana miRNA isolation kit, Thermo Fisher, Waltham, MA) from three replicates of 50 pooled ovaries for each condition in each genotype. In total, we prepared 18 samples (HP diapause, HP persistently reproductive, HP age-matched control, LP diapause, LP persistently reproductive, and LP age-matched control) and 18 libraries using NEBNext Ultra II (directional) with Poly-A selection, and sequenced libraries using Illumina 2×150 for a total of 30M reads per sample (Admera Health Biopharma Services, South Plainfield, NJ). All sequencing reads are available on the Sequencing Read Archive (NCBI), accession number PRJNA884433.

We trimmed raw reads using Trimmomatic (v.0.39) (89) and mapped reads to the *D. melanogaster* reference transcriptome using STAR aligner (v.2.7.10) (90). We estimated expression levels using FeatureCounts (v.2.0.3) (91) and analyzed differential expression using DESeq2 (v.1.36.0) in R (92). We discarded genes with fewer than 50 reads total across all samples in these analyses. We defined genes as significantly differentially expressed if the false discovery rate (FDR) was less than 0.05. Upon analyzing differential expression, we found that six and seven genes in the HP and LP lines, respectively, were significantly differentially expressed between diapausing and persistently reproductive ovaries (FDR < 0.05) but had a high standard error (lfcSE >1). In both genotypes, only a single replicate of the (pooled) persistently reproductive ovaries had an elevated number of reads mapping to these genes. We discovered that these genes belong to the multi-copy chorion gene cluster, which undergoes selective, 15-80-fold, gene amplification (endo-replication) in the ovarian follicle cells from ovary stages 8-14 (93). This observation is consistent with a small amount of contamination of a later stage egg chamber. Indeed, the chorion genes accounted for the deviation of the single replicate from the other two replicates in both genotypes. Furthermore, excluding these genes had no effect on the conclusions drawn from the data.

To compare diapause-specific gene expression between the HP and LP lines, we used the age-matched control ovaries to normalize genes that were differentially expressed between diapausing and persistently reproductive ovaries. We included in downstream analyses only those genes that were differentially expressed between diapausing and persistently reproductive ovaries *and* between diapausing and age-matched control ovaries in a given genotype. In other words, we excluded genes that are specifically up- or down-regulated in persistently reproductive ovaries compared to both diapause and age-matched controls. After generating this reduced list of significantly differentially expressed genes for both HP and LP genotypes, we compared across the two gene lists and determined which genes were differentially expressed in both HP and LP (“genotype-independent”), differentially expressed only in the HP line (“HP-specific”), or differentially expressed only in the LP line (“LP-specific”, see Fig. 4A).

We performed pathway enrichment analysis of differentially expressed genes using the Reactome Pathway Database (v.81) (94) We considered pathways significantly enriched if there were five or more genes in a given category and the FDR was less than 0.05.

### Western blotting and analysis

To assay histone mark abundance in the ovary, we isolated by microdissection ovary stages 1-7 (see above) in 1X PBS and ground the material in RIPA buffer (Cell Signaling Technology, Danvers, MA), Protease Inhibitor Cocktail (Roche, Basel, Switzerland), and PMSF (Cell Signaling Technology, Danvers, MA). To promote solubility, we incubated the lysate in Benzonase (Sigma Aldrich, St. Louis, MO) for 30 min at 4°C. We probed the blots with anti-H3K4me3 (Active Motif, Carlsbad, CA), anti-H3K36me1 (Abcam, Cambridge, UK), anti-H3K27me3 (Active Motif, Carlsbad, CA), anti-H3K27ac (Abcam, Cambridge, UK), anti-H3K9me3 (Abcam, Cambridge, UK), and anti-H3K9ac (Abcam, Cambridge, UK) at 1:1000 dilution. We probed with anti-α-tubulin (Developmental Studies Hybridoma Bank, Iowa City, IA) as a loading control (also 1:1000 dilution). We used anti-mouse and anti-rabbit HRP secondaries (Kindle Biosciences, Greenwich, CT) both at 1:1000. We exposed the blots with Kwikquant western blot detection kit and imaged with a Kwikquant imager (Kindle 277 Biosciences, Greenwich, CT). We quantified relative fluorescence of marks according to Stael *et al*. (2022) and normalized all measurements to diapause (95). We ran a third biological replicate only if we detected consistent differences in abundance across two replicates. For marks H3K4me3 and H3K36me1, we ran three biological replicates and compared abundance across diapausing and reproductive ovaries using Mann-Whitney U test.

### Tissue-specific knockdown of histone writers and erasers and analysis of diapause plasticity

To knockdown expression of histone mark writers and erasers, we took advantage of preconstructed D. melanogaster lines from the Transgenic RNAi Project (96). These lines encode “Upstream Activating Sequence” (UASp)-driven short hairpins (shRNA) that target transcripts encoding *D. melanogaster JHDM2* (Bloomington Drosophila Stock Center “BDSC” #32975), *Set2* (BDSC #55221), *Set1* (BDSC #33704), and *lid* (BDSC #36652). These cassettes are inserted into attP landing sites. Given the well-known effects of genetic background on diapause plasticity (42, 45, 66, 67), we carefully introgressed the chromosome encoding each UASp-shRNA construct (*Set2* and *lid* on chromosome II, *JHDM2* and *Set1* on chromosome III) into the HP inbred line (see Fig. S6 for crossing scheme). Furthermore, we ensured no recombination using a combination of balancer chromosomes and transmission only through males (which do not recombine) to tightly control the genetic background of the control and experimental flies (Fig. S6). Moreover, we only compared experimental genotypes encoding the UASp-shRNA construct in a given attP site to control genotypes encoding the same chromosome encoding the same attP site but lacking the UASp-shRNA construct (BDSC #36304 for chromosome II, BDSC #36303 for chromosome III). To verify the presence of these constructs after the multi-generation crossing scheme, we used PCR to amplify the AmpR gene introduced along with the UAS-shRNA construct (Table S1). We crossed these stable stocks to the MTD-Gal4 driver (BDSC #31777), which expresses the GAL4 transcription factor throughout the ovary (97).

To validate the knockdown of transcription by RNAi, we performed RT-qPCR on RNA prepared from ovaries stages 1-7 in control and experimental shRNA genotypes at 48 hours after eclosion at 25°C (Table S1). To validate the depletion or enrichment of histone mark abundance by RNAi against the target chromatin writers and erasers, we conducted western blotting (as above) on protein lysate prepared from stages 1-7 ovaries from control and experimental shRNA genotypes 48 hours after eclosion at 25°C. We note that RNAi efficiency decreases with decreasing temperature (98, 99), disabling us from distinguishing between the effects of RNAi on diapause entry and maintenance.

To determine whether experimental manipulation of histone mark abundance altered diapause plasticity, we assayed diapause in control and experimental RNAi lines by dissecting ovaries after 28 days in diapause-inducing conditions, as described above. We analyzed diapause plasticity using an odds ratio (implemented in R) comparing diapause plasticity between control and RNAi lines, which represents the change in likelihood of diapause given the presence of transcript knockdown.

*D. melanogaster* ovary development depends in part on chromatin-mediated gene regulation. We sought to rule out the possibility that the observed increase in incidence of arrest (“diapause plasticity”) upon transcript knockdown was simply due to a block in ovary development, independent of diapause. This was of particular concern given that experimental depletion of H3K36me1 and H3K4me3 increased diapause to nearly 100%. We reasoned that increasing the dynamic range of diapause plasticity in both experimental and control lines could allow us to determine if ovary development was blocked upon transcript knockdown at 12°C. To decrease the baseline diapause plasticity in both experimental and control lines, we took advantage of the transgenerational decrease in diapause plasticity (Fig. 1D). We exposed females from the UAS-shRNA and control lines to diapause conditions and assayed ovary development using the modified Bellymount method described above. We then crossed females in diapause to the MTD-Gal4 driver (BDSC #31777), and exposed daughters from this cross to simulated winter conditions and assayed for diapause plasticity. We found that fewer than 30% of these daughters have arrested ovaries, suggesting that *Set1* or *Set2* knockdown alone does not block ovary development at 12°C (Fig. S3B).

### Immunofluorescence and image analysis

To evaluate whether the LP line entered canonical diapause similarly to the temperate HP line, we conducted immunofluorescence on ovaries following (100). To assay germline stem cell number, we stained ovaries with anti-α-spectrin (1:300, Developmental Studies Hybridoma Bank, Iowa City, IA) and anti-Vasa (1:50, Developmental Studies Hybridoma Bank, Iowa City, IA). Following criteria described in (101), we defined germline stem cells as the anterior-most Vasa-positive cells in the stem cell niche that display an anterior α-spectrin signal (the “spectrosome”). Scanning through each z-stack, we counted the number of germline stem cells from 10 germaria in 5 ovary pairs, for a total of 50 germaria. We used a 2-way ANOVA to evaluate the statistical significance of the effects of treatment (diapause and age-matched control) and time (28 days in treatment vs. 5 days recovered from treatment). To assay double-strand break abundance, we stained ovaries with anti-γ-H2Av (1:1000, Developmental Studies Hybridoma Bank, Iowa City, IA). To quantify the average fluorescence of γ-H2Av in ovaries, we outlined representative stage 4 egg chambers with the Freehand tool in FIJI (v.1.0) (102). Also in FIJI, we calculated the fluorescent signal intensity using the polygon tool to define the borders of the tissue. We used the “measure tool” to calculate the mean pixels within these boundaries. We normalized all fluorescence intensity values of the HP line to the mean intensity value of the age-matched control in HP, and all fluorescence intensity values of the LP line to the mean intensity value of the age-matched control in LP. We calculated fluorescence from 10 replicates of stage four egg chambers in five ovary pairs, for a total of 50 egg chambers. We compared the mean fluorescence of γ-H2av in diapause and age-matched control ovaries using a Mann-Whitney U test (implemented in R).

For all immunofluorescence experiments, we mounted ovaries with ProLong Gold Antifade Reagent with DAPI (Thermo Fisher Scientific, Waltham, MA). We imaged slides at 63x magnification on a Leica TCS SP8 Four Channel Spectral Confocal System. For each experiment, we used the same imaging parameters across genotypes and reproductive state.

### Fertility assay

To further evaluate whether the LP line entered canonical diapause similarly to the temperate HP line, we assayed diapause-induced preservation of fertility in both lines. We counted progeny of females crossed to wildtype (w^1118^) males after diapause exit at 28 days in 12°C. We compared these females to age-matched controls (maintained at 25°C for 28 days) crossed in parallel to wildtype (w^1118^) males. In each vial, we crossed three females and six males. We replicated each cross across 12 vials and flipped each cross onto fresh food every three days. To exclude age-dependent male fertility effects, we replaced the six males every three days with one to three day old males. We recorded the number of adult progeny from each flip. We compared the mean number of progeny from diapause and age-matched control using a Mann-Whitney U test (implemented in R).

## ACKNOWLEDGEMENTS

We thank the Levine Lab, A. Das, D. Dudka, E. Joyce, D. Wagner, and N. Bonini for discussions about the project and the Levine Lab and D. Wagner for feedback on the manuscript. This work was supported by Start Up funds and University Research Fund award from the University of Pennsylvania School of Arts and Sciences to M.T.L and P.S.

## AUTHOR CONTRIBUTIONS

M.T.L., A.D.E., and P.S. designed the experiments. A.D.E. and R.A.F. performed the experiments. A.D.E. analyzed the experiments. M.T.L. and A.D.E. wrote the manuscript.

---

**Figure S1.**
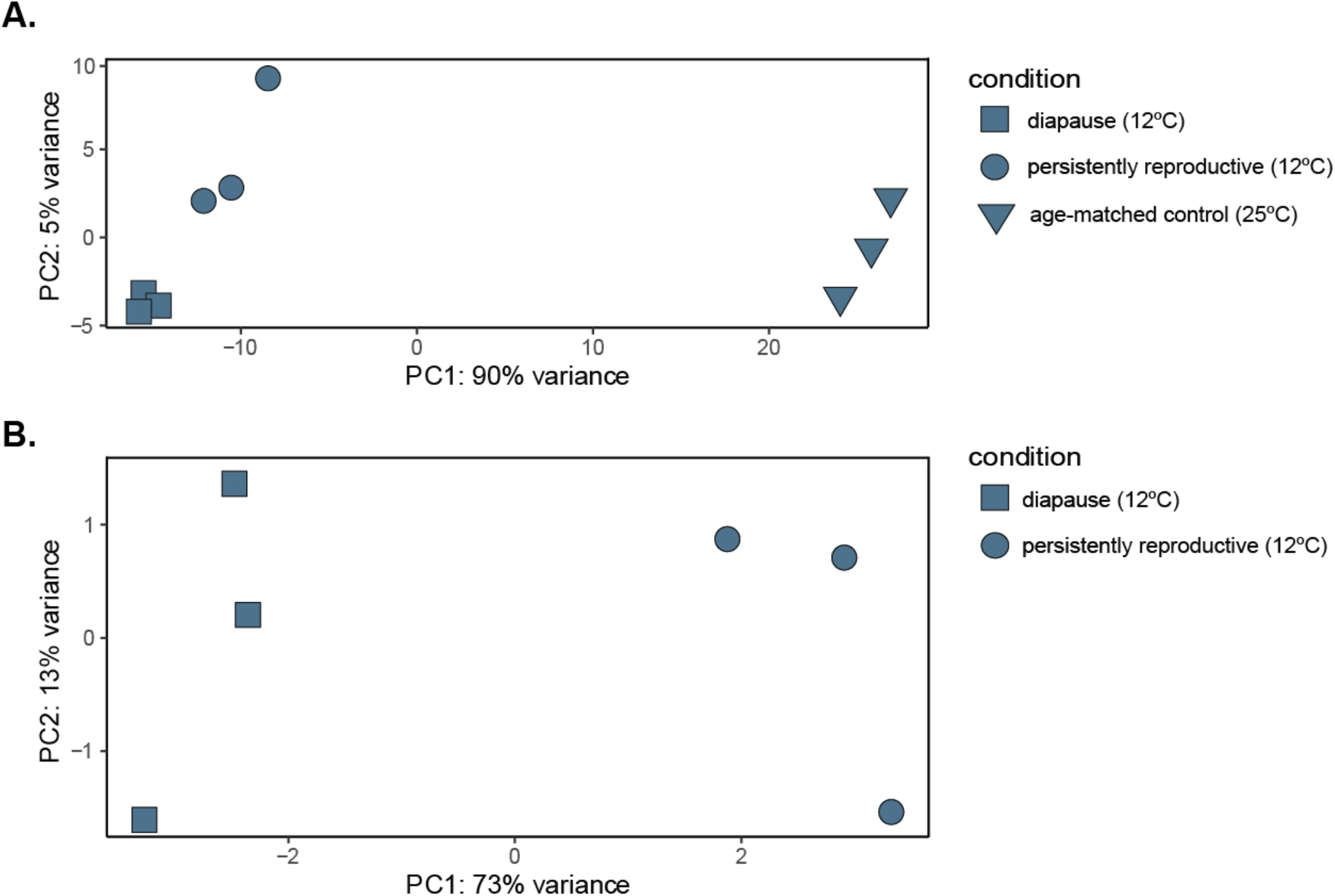
Principal component analysis (PCA) of RNA-seq reads from the temperate North American inbred line. (A) PCA of RNA-seq reads from diapausing (square), persistently reproductive (circle), and age-matched control (triangle) ovaries, stages 1-7 only. Note that temperature explains most of the variance between the three samples (PC1, 90%). (B) PCA of RNA-seq reads from diapausing and persistently reproductive ovaries only.

**Figure S2.**
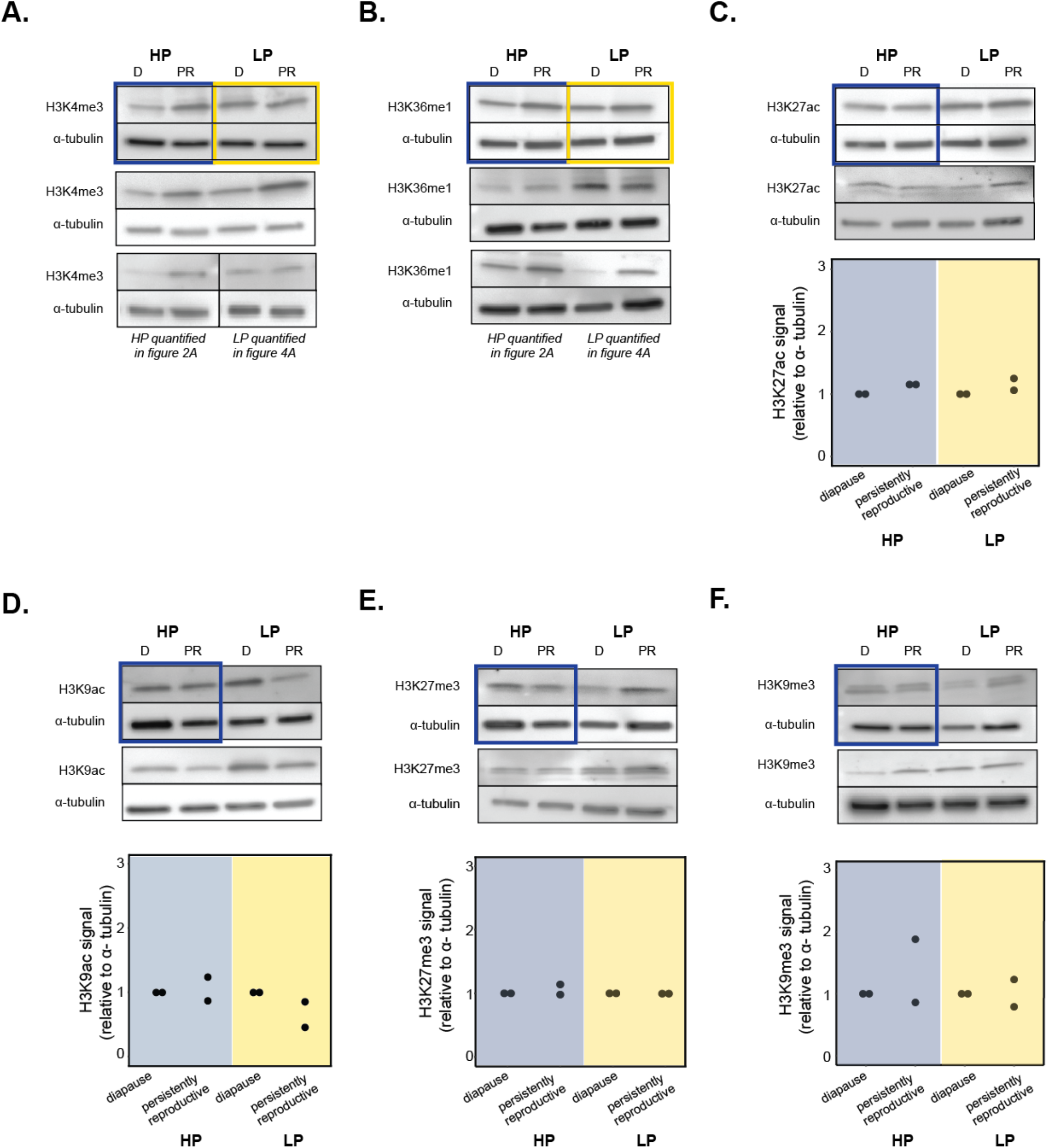
Western blots probed for various histone marks on multiple biological replicates. Blots of ovary lysate prepared from diapausing and persistently reproductive ovaries and quantification relative to a-tubulin loading control of (A) H3K4me3, (B) H3K36me1, (C) H3K27ac, (D) H3K9ac, (E) H3K27me3, and (F) H3K9me3. Blue boxes delineate replicates shown in Fig. 2A, yellow boxes delineate replicates shown in Fig. 4B. D = diapause, PR = persistently reproductive.

**Figure S3.**
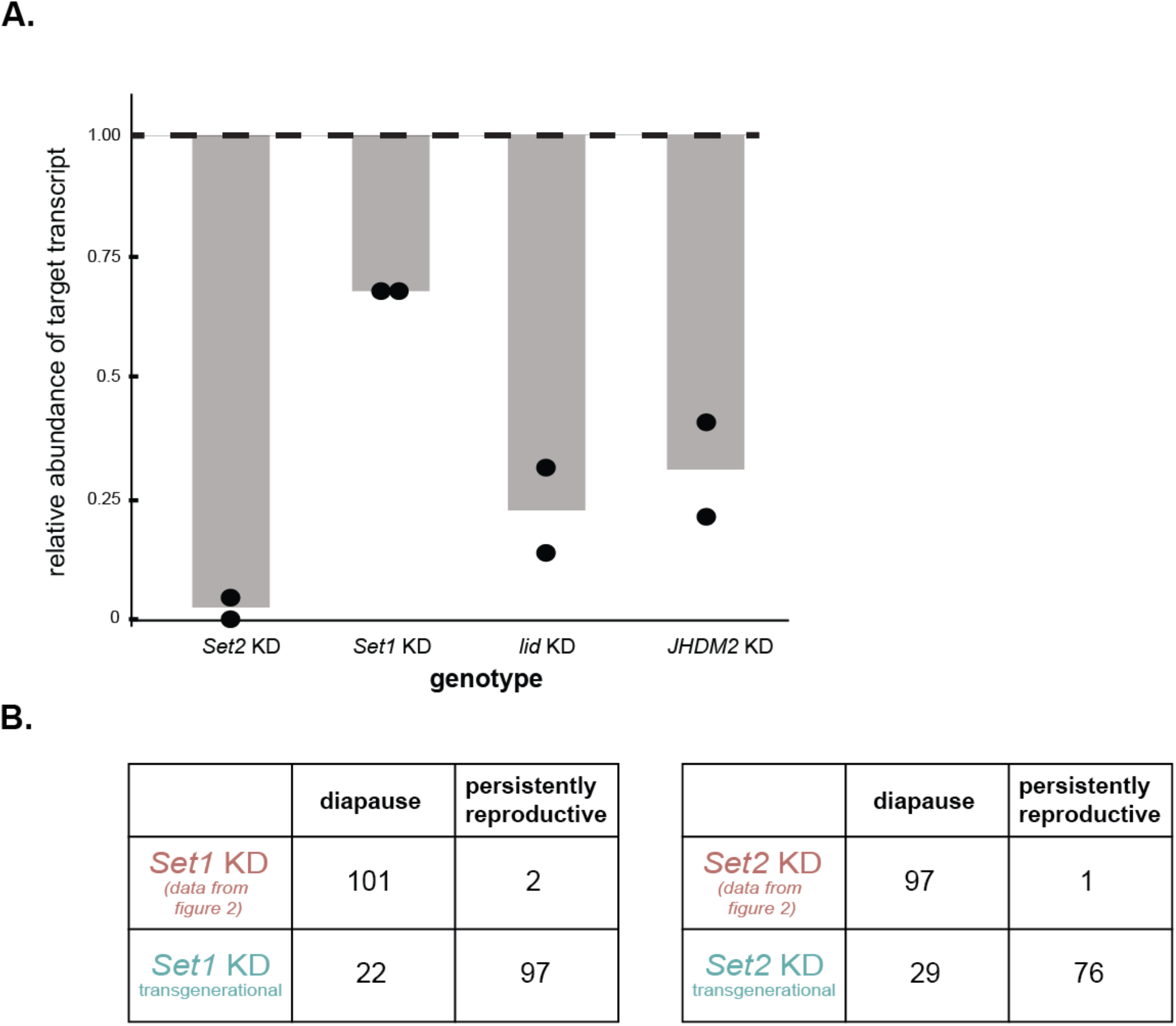
Quality control of RNAi experiments. (A) RT-qPCR confirming knockdown (KD) of transcripts targeted by RNAi. Note *Set2* and *lid* knockdown genotypes are compared to chromosome II control genotype, while *Set1* and *JHDM2* knockdown genotypes are compared to chromosome III control genotype. (B) Comparison of diapause plasticity upon histone writer knockdown in the ovaries of females whose mothers that had either undergone diapause (“transgenerational”) or not (data from Fig. 2). The abundance of persistently reproductive ovaries in both genotypes under the transgenerational treatment verified that knockdown of *Set1* or *Set2* alone does not block ovary development at 12°C (see Methods). “chr.” = chromosome.

**Figure S4.**
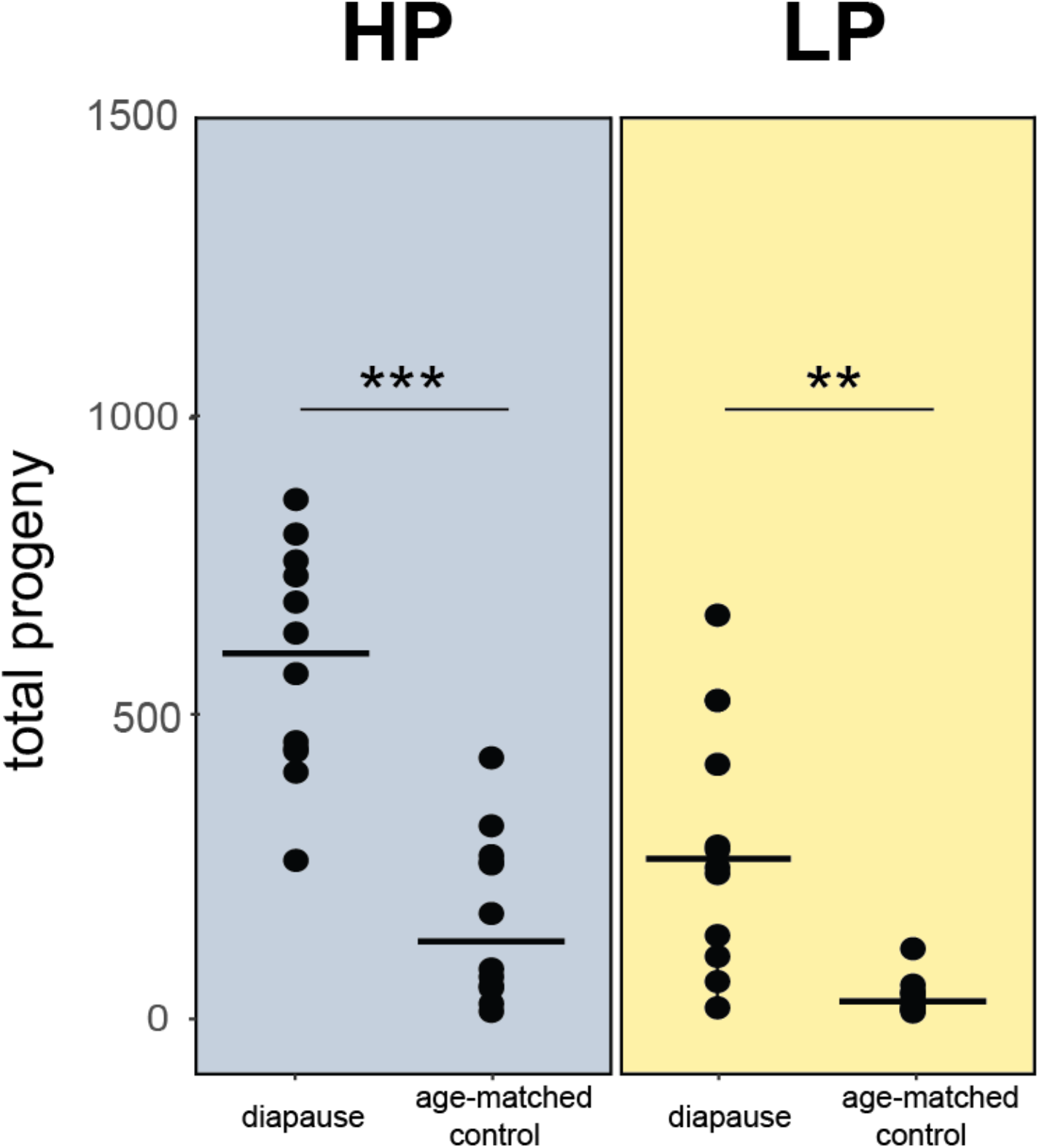
Total progeny from diapause and age-matched control females of HP (blue) and LP line (yellow). Each replicate represents a vial of three females. t-test, *** p<0.001, ** p<0.01

**Figure S5.**
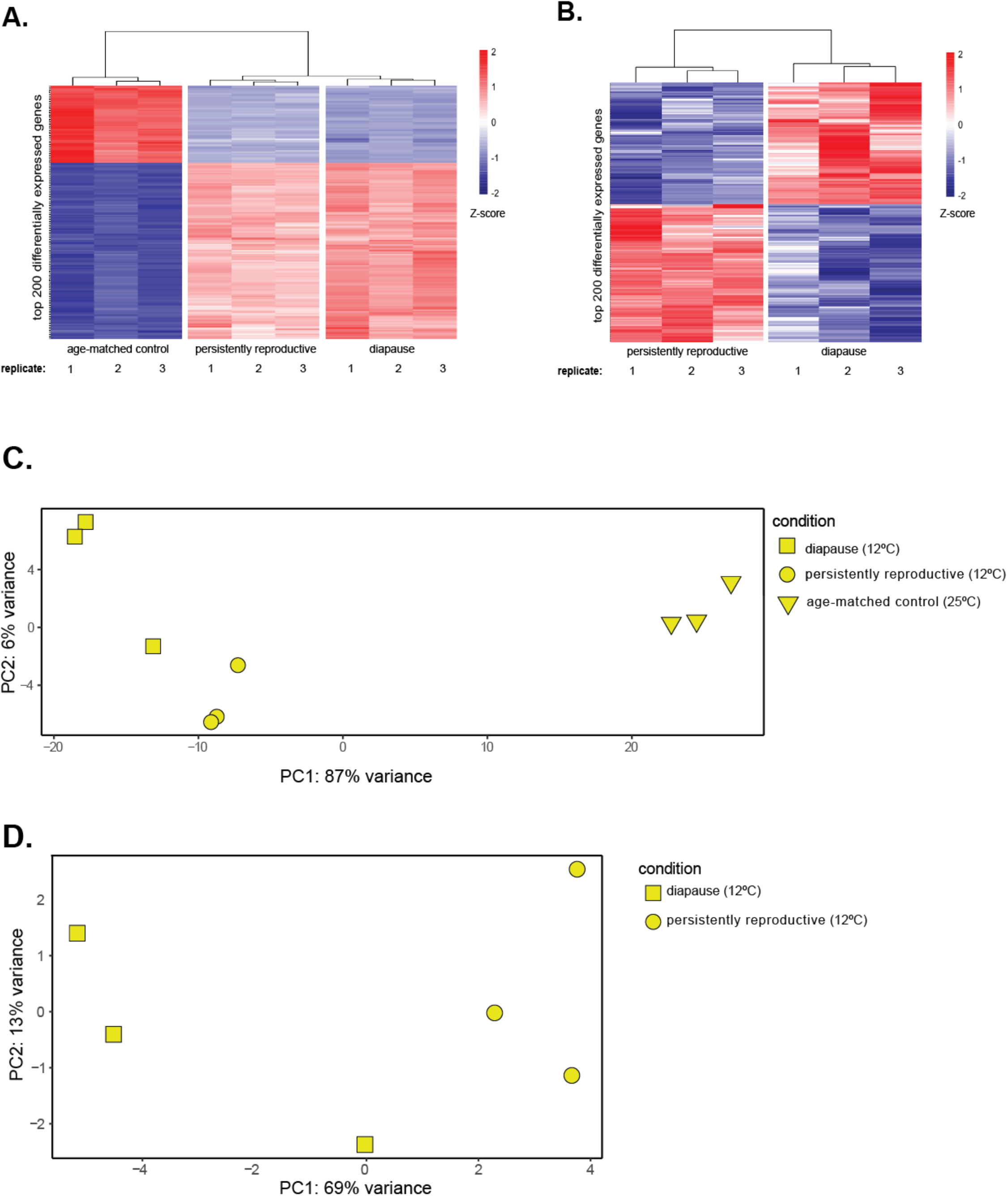
Heat map and principal component analysis (PCA) of LP line RNA-seq reads. (A) Heatmap of the top 200 significantly differentially expressed genes (by FDR) between age-matched control, diapausing, and persistently reproductive ovaries, stages 1-7 only. Blue-red gradient depicts the Z-score of each gene. Red corresponds to upregulated genes and blue corresponds to downregulated genes. (B) Heatmap of the top 200 significantly differentially expressed genes (by FDR) between diapausing and persistently reproductive ovaries only. Blue-red gradient depicts the Z-score of each gene. Red corresponds to upregulated genes and blue corresponds to downregulated genes. (C) PCA of RNA-seq reads from diapausing (square), persistently reproductive (circle), and age-matched control (triangle) ovaries. Note that temperature explains most of the variance among the three samples (PC1, 87%). (D) PCA of RNA-seq reads from diapausing and persistently reproductive ovaries.

**Figure S6.**
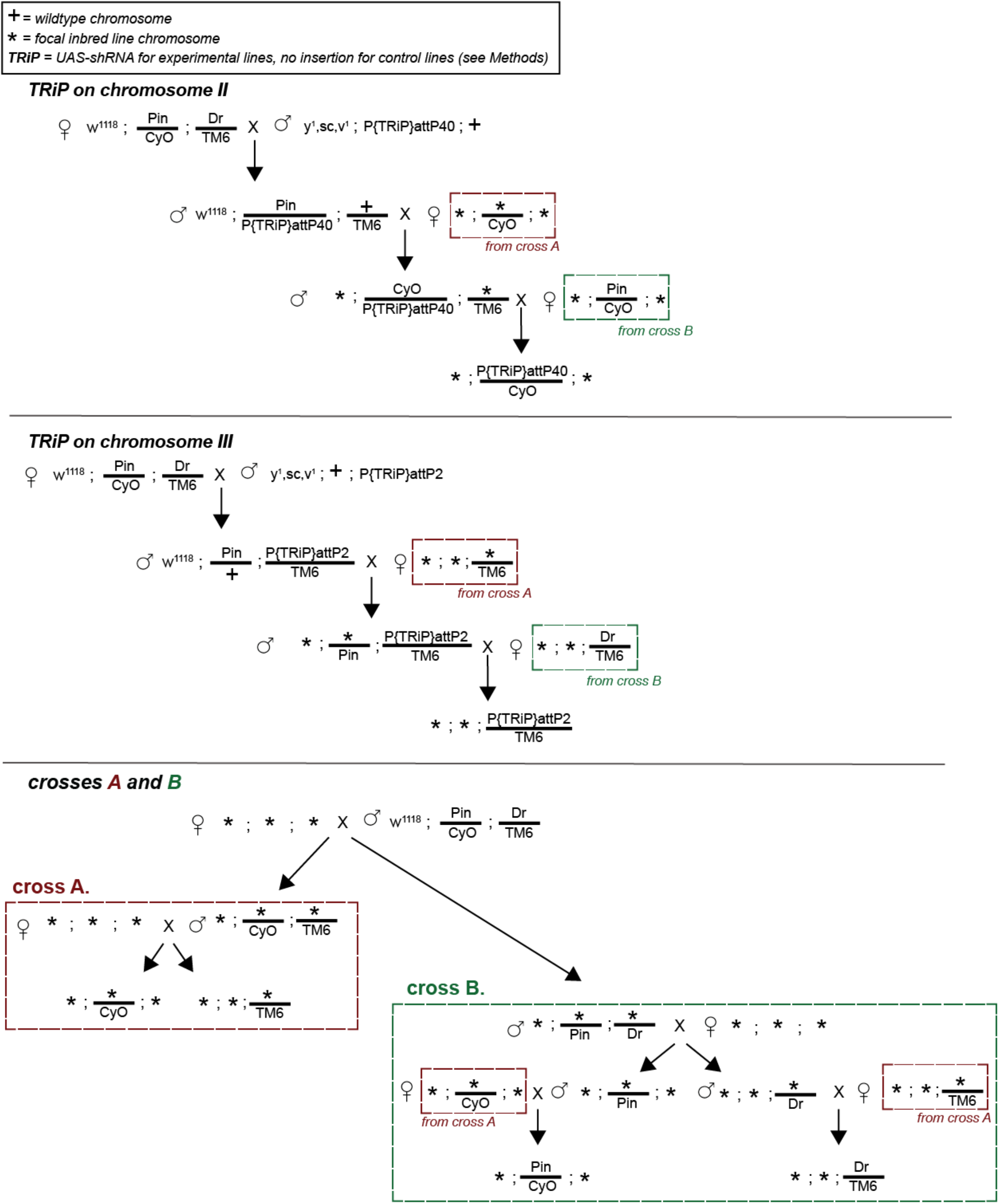
Crossing scheme used to generate RNAi and control lines. Brown dashed boxes correspond to lines constructed from cross A (bottom) and green dashed boxes correspond to lines constructed from cross B (bottom).

## SUPPLEMENTARY TABLE LEGEND

**Table S1.**
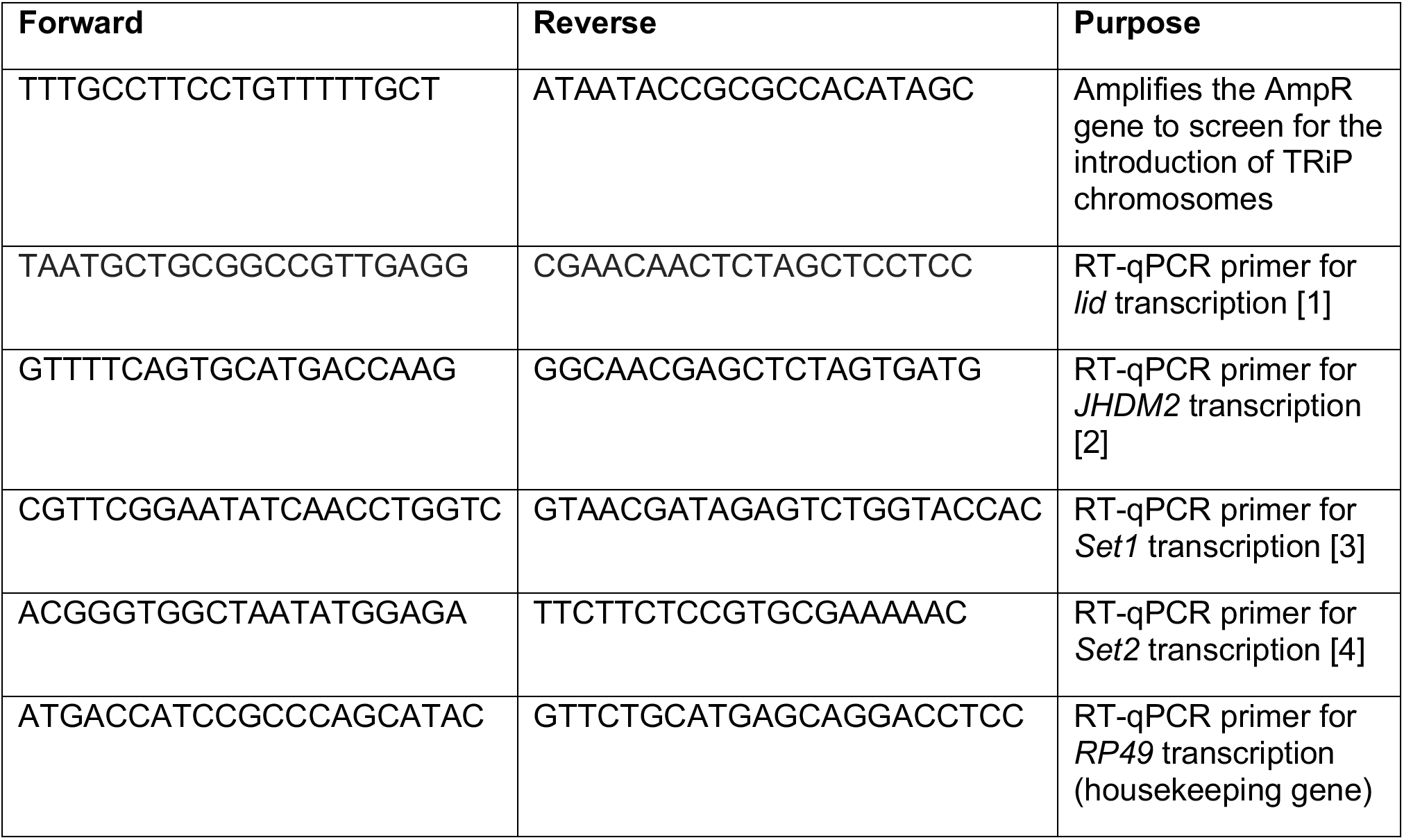
Primers used in study.

## LITERATURE CITED

1. Mills LS, Zimova M, Oyler J, Running S, Abatzoglou JT, Lukacs PM. Camouflage mismatch in seasonal coat color due to decreased snow duration. Proceedings of the National Acadademy of Sciences. 2013;110(18):7360–5.

2. Fielenbach N, Antebi A. *C. elegans* dauer formation and the molecular basis of plasticity. Genes & Development. 2008;22(16):2149–65.

3. Cassada RC, Russell RL. The dauerlarva, a post-embryonic developmental variant of the nematode *Caenorhabditis elegans*. Developmental Biology. 1975;46(2):326–42.

4. Sommer RJ. Phenotypic plasticity: from theory and genetics to current and future challenges. Genetics. 2020;215(1):1–13.

5. Beldade P, Mateus AR, Keller RA. Evolution and molecular mechanisms of adaptive developmental plasticity. Molecular Ecology. 2011;20(7):1347–63.

6. Lafuente E, Beldade P. Genomics of developmental plasticity in animals. Frontiers in Genetics. 2019;10:720.

7. Weiss LC. Sensory ecology of predator-induced phenotypic plasticity. Frontiers in Behavioral Neuroscience. 2018;12:330.

8. Yang CH, Andrew Pospisilik J. Polyphenism - a window into gene-environment interactions and phenotypic plasticity. Frontiers in Genetics. 2019;10:132.

9. Turner BM. Cellular memory and the histone code. Cell. 2002;111(3):285–91.

10. Papp B, Plath K. Epigenetics of reprogramming to induced pluripotency. Cell. 2013;152(6):1324–43.

11. Ringrose L, Paro R. Polycomb/Trithorax response elements and epigenetic memory of cell identity. Development. 2007;134(2):223–32.

12. Lee N, Maurange C, Ringrose L, Paro R. Suppression of Polycomb group proteins by JNK signalling induces transdetermination in *Drosophila* imaginal discs. Nature. 2005;438(7065):234–7.

13. Chen T, Dent SY. Chromatin modifiers and remodellers: regulators of cellular differentiation. Nature Reviews Genetics. 2014;15(2):93–106.

14. Smith ZD, Meissner A. DNA methylation: roles in mammalian development. Nature Reviews Genetics. 2013;14(3):204–20.

15. Skene PJ, Henikoff S. Histone variants in pluripotency and disease. Development. 2013;140(12):2513–24.

16. Chen X, Hiller M, Sancak Y, Fuller MT. Tissue-specific TAFs counteract Polycomb to turn on terminal differentiation. Science. 2005;310(5749):869–72.

17. Lee LR, Wengier DL, Bergmann DC. Cell-type-specific transcriptome and histone modification dynamics during cellular reprogramming in the *Arabidopsis* stomatal lineage. Proceedings of the National Academy of Sciences. 2019;116(43):21914–24.

18. Valk-Lingbeek ME, Bruggeman SW, van Lohuizen M. Stem cells and cancer; the polycomb connection. Cell. 2004;118(4):409–18.

19. Spivakov M, Fisher AG. Epigenetic signatures of stem-cell identity. Nature Reviews Genetics. 2007;8(4):263–71.

20. Ashburner M. Patterns of puffing activity in the salivary gland chromosomes of *Drosophila*. I. Autosomal puffing patterns in a laboratory stock of *Drosophila melanogaster*. Chromosoma. 1967;21(4):398–428.

21. Ashburner M. Patterns of puffing activity in the salivary gland chromosomes of *Drosophila*. V. Responses to environmental treatments. Chromosoma. 1970;31(3):356–76.

22. Berthiaume M, Boufaied N, Moisan A, Gaudreau L. High levels of oxidative stress globally inhibit gene transcription and histone acetylation. DNA and Cell Biology. 2006;25(2):124–34.

23. Causton HC, Ren B, Koh SS, Harbison CT, Kanin E, Jennings EG, et al. Remodeling of yeast genome expression in response to environmental changes. Molecular Biology of the Cell. 2001;12(2):323–37.

24. Uffenbeck SR, Krebs JE. The role of chromatin structure in regulating stress-induced transcription in *Saccharomyces cerevisiae*. Biochemistry and Cell Biology. 2006;84(4):477–89.

25. Gowen JW, Gay EH. Chromosome constitution and behavior in eversporting and mottling in *Drosophila melanogaster*. Genetics. 1934;19(3):189–208.

26. Zhao J, Herrera-Diaz J, Gross DS. Domain-wide displacement of histones by activated heat shock factor occurs independently of Swi/Snf and is not correlated with RNA polymerase II density. Molecular and Cellular Biology. 2005;25(20):8985–99.

27. Kim JM, Sasaki T, Ueda M, Sako K, Seki M. Chromatin changes in response to drought, salinity, heat, and cold stresses in plants. Frontiers in Plant Science. 2015;6:114.

28. Dai H, Wang Z. Histone modification patterns and their responses to environment. Current Environmental Health Reports. 2014;1(1):11–21.

29. Ozawa T, Mizuhara T, Arata M, Shimada M, Niimi T, Okada K, et al. Histone deacetylases control module-specific phenotypic plasticity in beetle weapons. Proceedings of the National Academy of Sciences. 2016;113(52):15042–7.

30. Simola DF, Graham RJ, Brady CM, Enzmann BL, Desplan C, Ray A, et al. Epigenetic (re)programming of caste-specific behavior in the ant *Camponotus floridanus*. Science. 2016;351(6268).

31. Kucharski R, Maleszka J, Foret S, Maleszka R. Nutritional control of reproductive status in honeybees via DNA methylation. Science. 2008;319(5871):1827–30.

32. Hu C-K, Wang W, Brind’Amour J, Singh PP, Reeves GA, Lorincz MC, et al. Vertebrate diapause preserves organisms long term through Polycomb complex members. Science. 2020;367(6480):870–4.

33. Lu YX, Denlinger DL, Xu WH. Polycomb repressive complex 2 (PRC2) protein ESC regulates insect developmental timing by mediating H3K27me3 and activating prothoracicotropic hormone gene expression. Journal of Biological Chemistry. 2013;288(32):23554–64.

34. Pegoraro M, Bafna A, Davies NJ, Shuker DM, Tauber E. DNA methylation changes induced by long and short photoperiods in *Nasonia*. Genome Research. 2016;26(2):203–10.

35. Herman JJ, Sultan SE. DNA methylation mediates genetic variation for adaptive transgenerational plasticity. Proceedings of the Royal Society B: Biological Sciences. 2016;283(1838).

36. Saunders D, Richard D, Applebaum S, Ma M, Gilbert L. Photoperiodic diapause in *Drosophila melanogaster* involves a block to the juvenile hormone regulation of ovarian maturation. General and comparative endocrinology. 1990;79(2):174–84.

37. Kubrak OI, Kucerova L, Theopold U, Nassel DR. The Sleeping Beauty: How Reproductive Diapause Affects Hormone Signaling, Metabolism, Immune Response and Somatic Maintenance in *Drosophila melanogaster*. Plos One. 2014;9(11):e113051.

38. Kučerová L, Kubrak OI, Bengtsson JM, Strnad H, Nylin S, Theopold U, et al. Slowed aging during reproductive dormancy is reflected in genome-wide transcriptome changes in *Drosophila melanogaster*. BMC Genomics. 2016;17(1).

39. Allen M. What makes a fly enter diapause? Fly. 2007;1(6):307–10.

40. Koštál V. Eco-physiological phases of insect diapause. Journal of Insect Physiology. 2006;52(2):113–27.

41. Saunders DS, Henrich VC, Gilbert LI. Induction of diapause in *Drosophila melanogaster*: photoperiodic regulation and the impact of arrhythmic clock mutations on time measurement. Proceedings of the National Academy of Sciences. 1989;86(10):3748–52.

42. Saunders DS, Gilbert LI. Regulation of ovarian diapause in *Drosophila melanogaster* by photoperiod and moderately low temperature. Journal of Insect Physiology. 1990;36(3):195–200.

43. Tatar M, Chien Susan A, Priest Nicholas K. Negligible senescence during reproductive dormancy in *Drosophila melanogaster*. The American Naturalist. 2001;158(3):248–58.

44. Schmidt PS, Matzkin L, Ippolito M, Eanes WF. Geographic variation in diapause incidence, life-history traits, and climatic adaptation in *Drosophila melanogaster*. Evolution. 2005;59(8):1721–32.

45. Schmidt PS, Paaby AB, Heschel MS. Genetic variance for diapause expression and associated life histories in *Drosophila melanogaster*. Evolution. 2005;59(12).

46. Zhao X, Bergland AO, Behrman EL, Gregory BD, Petrov DA, Schmidt PS. Global transcriptional profiling of diapause and climatic adaptation in *Drosophila melanogaster*. Molecular Biology and Evolution. 2016;33(3):707–20.

47. Tatar M, Yin C. Slow aging during insect reproductive diapause: why butterflies, grasshoppers and flies are like worms. Experimental Gerontology. 2001;36(4-6):723–38.

48. Easwaran S, Van Ligten M, Kui M, Montell DJ. Enhanced germline stem cell longevity in *Drosophila* diapause. Nature Communications. 2022;13(1).

49. Erickson PA, Weller CA, Song DY, Bangerter AS, Schmidt P, Bergland AO. Unique genetic signatures of local adaptation over space and time for diapause, an ecologically relevant complex trait, in *Drosophila melanogaster*. PLoS Genetics. 2020;16(11).

50. Schmidt PS, Paaby AB. Reproductive diapause and life-history clines in North American populations of *Drosophila melanogaster*. Evolution. 2008;62(5):1204–15.

51. Schmidt PS, Conde DR. Environmental heterogeneity and the maintenance of genetic variation for reproductive diapause in *Drosophila melanogaster*. Evolution. 2006;60(8):1602–11.

52. Anway MD, Cupp AS, Uzumcu M, Skinner MK. Epigenetic transgenerational actions of endocrine disruptors and male fertility. Science. 2005;308(5727):1466–9.

53. Newbold RR, Padilla-Banks E, Jefferson WN. Adverse effects of the model environmental estrogen diethylstilbestrol are transmitted to subsequent generations. Endocrinology. 2006;147(6):s11–s7.

54. Nilsson EE, Skinner MK. Environmentally induced epigenetic transgenerational inheritance of reproductive disease. Biology of Reproduction. 2015;93(6).

55. Casier K, Boivin A, Carré C, Teysset L. Environmentally-induced transgenerational epigenetic inheritance: implication of PIWI interacting RNAs. Cells. 2019;8(9).

56. Houri-Ze’evi L, Korem Y, Sheftel H, Faigenbloom L, Toker IA, Dagan Y, et al. A tunable mechanism determines the duration of the transgenerational small RNA inheritance in *C. elegans*. Cell. 2016;165(1):88–99.

57. Klosin A, Lehner B. Mechanisms, timescales and principles of trans-generational epigenetic inheritance in animals. Current Opinion in Genetics & Development. 2016;36:41–9.

58. Jaenisch R, Bird A. Epigenetic regulation of gene expression: how the genome integrates intrinsic and environmental signals. Nature Genetics. 2003;33(S3):245–54.

59. Levine MT, Eckert ML, Begun DJ. Whole-Genome Expression Plasticity across Tropical and Temperate *Drosophila melanogaster* Populations from Eastern Australia. Molecular Biology and Evolution. 2010;28(1):249–56.

60. Huang W, Carbone MA, Lyman RF, Anholt RRH, Mackay TFC. Genotype by environment interaction for gene expression in *Drosophila melanogaster*. Nature Communications. 2020;11(1).

61. Quina AS, Buschbeck M, Di Croce L. Chromatin structure and epigenetics. Biochemical Pharmacology. 2006;72(11):1563–9.

62. Oudet P, Gross-Bellard M, Chambon P. Electron microscopic and biochemical evidence that chromatin structure is a repeating unit. Cell. 1975;4(4):281–300.

63. Kouzarides T. Chromatin modifications and their function. Cell. 2007;128(4):693–705.

64. Lawrence M, Daujat S, Schneider R. Lateral thinking: how histone modifications regulate gene expression. Trends in Genetics. 2016;32(1):42–56.

65. Zhang T, Cooper S, Brockdorff N. The interplay of histone modifications - writers that read. EMBO Reports. 2015;16(11):1467–81.

66. Williams KD, Sokolowski MB. Diapause in *Drosophila melanogaster* females: a genetic analysis. Heredity. 1993;71(3):312–7.

67. Emerson KJ, Uyemura AM, McDaniel KL, Schmidt PS, Bradshaw WE, Holzapfel CM. Environmental control of ovarian dormancy in natural populations of *Drosophila melanogaster*. Journal of Comparative Physiology A. 2009;195(9):825–9.

68. Lirakis M, Dolezal M, Schlötterer C. Redefining reproductive dormancy in *Drosophila* as a general stress response to cold temperatures. Journal of Insect Physiology. 2018;107:175–85.

69. Kacsoh BZ, Bozler J, Ramaswami M, Bosco G. Social communication of predator-induced changes in *Drosophila* behavior and germ line physiology. eLife. 2015;4.

70. Serizier SB, McCall K. Scrambled eggs: apoptotic cell clearance by non-professional phagocytes in the *Drosophila* ovary. Frontiers in Immunology. 2017;8.

71. Drummond-Barbosa D, Spradling AC. Stem cells and their progeny respond to nutritional changes during *Drosophila* oogenesis. Developmental biology. 2001;231(1):265–78.

72. Buszczak M, Cooley L. Eggs to die for: cell death during *Drosophila* oogenesis. Cell Death & Differentiation. 2000;7(11): 1071–4.

73. Skene PJ, Henikoff S. An efficient targeted nuclease strategy for high-resolution mapping of DNA binding sites. eLife. 2017;6.

74. Lyko F, Ramsahoye BH, Jaenisch R. DNA methylation in *Drosophila melanogaster*. Nature. 2000;408(6812):538–40.

75. Kassis JA, Brown JL. Polycomb Group Response Elements in *Drosophila* and Vertebrates. Advances in Genetics2013. p. 83–118.

76. Booth Lauren N, Brunet A. The aging epigenome. Molecular Cell. 2016;62(5):728–44.

77. Li L, Greer C, Eisenman RN, Secombe J. Essential functions of the histone demethylase lid. PLoS Genetics. 2010;6(11):e1001221.

78. Chamani IJ, Keefe DL. Epigenetics and female reproductive aging. Frontiers in Endocrinology. 2019;10.

79. Pal S, Tyler JK. Epigenetics and aging. Science Advances. 2016;2(7).

80. Kaufman PD, Rando OJ. Chromatin as a potential carrier of heritable information. Current Opinion in Cell Biology. 2010;22(3):284–90.

81. Sarkies P. Molecular mechanisms of epigenetic inheritance: Possible evolutionary implications. Seminars in Cell & Developmental Biology. 2020;97:106–15.

82. Reynolds JA, Clark J, Diakoff SJ, Denlinger DL. Transcriptional evidence for small RNA regulation of pupal diapause in the flesh fly, *Sarcophaga bullata*. Insect Biochemistry and Molecular Biology. 2013;43(10):982–9.

83. Webster AK, Jordan JM, Hibshman JD, Chitrakar R, Baugh LR. Transgenerational effects of extended dauer diapause on starvation survival and gene expression plasticity in *Caenorhabditis elegans*. Genetics. 2018;210(1):263–74.

84. Machado HE, Bergland AO, O’Brien KR, Behrman EL, Schmidt PS, Petrov DA. Comparative population genomics of latitudinal variation in *Drosophila simulans* and *Drosophila melanogaster*. Molecular Ecology. 2016;25(3):723–40.

85. Levine MT, Begun DJ. Evidence of spatially varying selection acting on four chromatin-remodeling loci in *Drosophila melanogaster*. Genetics. 2008;179(1):475–85.

86. Reinhardt JA, Kolaczkowski B, Jones CD, Begun DJ, Kern AD. Parallel geographic variation in *Drosophila melanogaster*. Genetics. 2014;197(1):361–73.

87. Wuebbles DJ, Fahey DW, Hibbard KA, Dokken DJ, Stewart BC, Maycock TK. Climate science special report: fourth national climate assessment. US Global Change Research Program. 2017;1.

88. Edgar B, Koyama LAJ, Aranda-Díaz A, Su Y-H, Balachandra S, Martin JL, et al. Bellymount enables longitudinal, intravital imaging of abdominal organs and the gut microbiota in adult *Drosophila*. PLoS Biology. 2020;18(1).

89. Bolger AM, Lohse M, Usadel B. Trimmomatic: a flexible trimmer for Illumina sequence data. Bioinformatics. 2014;30(15):2114–20.

90. Dobin A, Davis CA, Schlesinger F, Drenkow J, Zaleski C, Jha S, et al. STAR: ultrafast universal RNA-seq aligner. Bioinformatics. 2013;29(1):15–21.

91. Liao Y, Smyth GK, Shi W. featureCounts: an efficient general purpose program for assigning sequence reads to genomic features. Bioinformatics. 2013;30(7):923–30.

92. Love MI, Huber W, Anders S. Moderated estimation of fold change and dispersion for RNA-seq data with DESeq2. Genome Biology. 2014;15(12).

93. Orr-Weaver TL. *Drosophila* chorion genes: cracking the eggshell’s secrets. BioEssays. 1991;13(3):97–105.

94. Gillespie M, Jassal B, Stephan R, Milacic M, Rothfels K, Senff-Ribeiro A, et al. The reactome pathway knowledgebase 2022. Nucleic Acids Research. 2022;50(D1):D687–D92.

95. Stael S, Miller LP, Fernández-Fernández ÁD, Van Breusegem F. Detection of damage-activated metacaspase activity by western blot in plants. Plant Proteases and Plant Cell Death. Methods in Molecular Biology2022. p. 127–37.

96. Perkins LA, Holderbaum L, Tao R, Hu Y, Sopko R, McCall K, et al. The Transgenic RNAi Project at Harvard Medical School: resources and validation. Genetics. 2015;201(3):843–52.

97. Petrella LN, Smith-Leiker T, Cooley L. The Ovhts polyprotein is cleaved to produce fusome and ring canal proteins required for *Drosophila* oogenesis. Development. 2007;134(4):703–12.

98. Szittya G. Low temperature inhibits RNA silencing-mediated defence by the control of siRNA generation. The EMBO Journal. 2003;22(3):633–40.

99. Brand AH, Manoukian AS, Perrimon N. Chapter 33: Ectopic expression in *Drosophila*. Methods in Cell Biology. Methods in Cell Biology1994. p. 635–54.

100. McKim KS, Joyce EF, Jang JK. Cytological analysis of meiosis in fixed *Drosophila* ovaries. Meiosis. Methods in Molecular Biology2009. p. 197–216.

101. Xie T, Spradling AC. A niche maintaining germ line stem cells in the *Drosophila* ovary. Science. 2000;290(5490):328–30.

102. Schindelin J, Arganda-Carreras I, Frise E, Kaynig V, Longair M, Pietzsch T, et al. Fiji: an open-source platform for biological-image analysis. Nature Methods. 2012;9(7):676–82.

## SUPPLEMENTARY REFERENCES

1. Drelon, C., H.M. Belalcazar, and J. Secombe, *The Histone Demethylase KDM5 Is Essential for Larval Growth in Drosophila*. Genetics, 2018. 209(3): p. 773–787.

2. Lorbeck, M.T., N. Singh, A. Zervos, M. Dhatta, M. Lapchenko, et al., The histone demethylase Dmel\Kdm4A controls genes required for life span and male-specific sex determination in Drosophila. Gene, 2010. 450(1-2): p. 8–17.

3. Hallson, G., R.E. Hollebakken, T. Li, M. Syrzycka, I. Kim, et al., dSet1 Is the Main H3K4 Di-and Tri-Methyltransferase Throughout Drosophila Development. Genetics, 2012. 190(1): p. 91–100.

4. Stabell, M., J. Larsson, R.B. Aalen, and A. Lambertsson, Drosophila dSet2 functions in H3-K36 methylation and is required for development. Biochemical and Biophysical

